# Identifying a gene signature of metastatic potential by linking pre-metastatic state to ultimate metastatic fate

**DOI:** 10.1101/2024.08.14.607813

**Authors:** Jesse S Handler, Zijie Li, Rachel K Dveirin, Weixiang Fang, Hani Goodarzi, Elana J Fertig, Reza Kalhor

## Abstract

Identifying the key molecular pathways that enable metastasis by analyzing the eventual metastatic tumor is challenging because the state of the founder subclone likely changes following metastatic colonization. To address this challenge, we labeled primary mouse pancreatic ductal adenocarcinoma (PDAC) subclones with DNA barcodes to characterize their pre-metastatic state using ATAC-seq and RNA-seq and determine their relative *in vivo* metastatic potential prospectively. We identified a gene signature separating metastasis-high and metastasis-low subclones orthogonal to the normal-to-PDAC and classical-to-basal axes. The metastasis-high subclones feature activation of IL-1 pathway genes and high NF-κB and Zeb/Snail family activity and the metastasis-low subclones feature activation of neuroendocrine, motility, and Wnt pathway genes and high CDX2 and HOXA13 activity. In a functional screen, we validated novel mediators of PDAC metastasis in the IL-1 pathway, including the NF-κB targets *Fos* and *Il23a*, and beyond the IL-1 pathway including *Myo1b* and *Tmem40*. We scored human PDAC tumors for our signature of metastatic potential from mouse and found that metastases have higher scores than primary tumors. Moreover, primary tumors with higher scores are associated with worse prognosis. We also found that our metastatic potential signature is enriched in other human carcinomas, suggesting that it is conserved across epithelial malignancies. This work establishes a strategy for linking cancer cell state to future behavior, reveals novel functional regulators of PDAC metastasis, and establishes a method for scoring human carcinomas based on metastatic potential.

## Introduction

Most patients with solid malignancies die from complications of metastasis^1^. This is true across diverse cancer subtypes, with few exceptions (e.g., glioblastoma). Due to the major clinical burden of metastasis, there is an urgent need to develop therapies targeting the metastatic cascade. However, we are currently hindered in our ability to develop such therapies by an incomplete understanding of the molecular mechanisms underlying metastasis.

Lineage tracing studies utilizing patient samples have demonstrated that across multiple cancer subtypes, metastases develop from a small number of subclones within heterogeneous primary tumors^2–6^, indicating the existence of subclones with an intrinsic advantage in progressing through the metastatic cascade. The exact nature of this advantage is unclear – genetic comparison of metastases and primary tumors in patient samples has not identified clear individual drivers of metastasis^7–9^, suggesting that the metastatic advantage is conferred by small contributions from a large set of factors. However, identifying these factors by analyzing the eventual metastatic tumor is challenging because the massive expansion of cell numbers after the completion of the metastatic cascade, which coincides with intense cell-cell competition and selection within the tumor, is likely to diminish or mask the features that conferred the initial advantage in completing the metastatic cascade. Therefore, the observed cell states and active molecular pathways in a metastatic tumor are likely different than those that enabled the founder of that metastatic tumor to successfully progress through the cascade.

Prior studies have sought to address this barrier by using somatic mutations^10,11^ or engineered DNA barcodes^12–14^ to identify the primary tumor subclones most closely related to the metastatic tumor and then characterize the molecular state of these subclones. These approaches have described transcriptomic features associated with metastatic potential, identifying a hybrid EMT (epithelial-to-mesenchymal transition) state in metastatic subclones in multiple cancer subtypes. However, a key limitation in this type of analysis is that the cell states of such retrospectively-defined primary tumor subclones that share a common ancestor with metastases are likely to be different from the cells that founded the metastases because of cancer cells’ transcriptional plasticity and the changing selective pressures in the growing primary tumor over time. Other studies have labeled subclones to prospectively to compare their metastatic potential^15,16^. However, these studies either characterized a number of subclones too small to be representative or used immunocompromised hosts which precludes interrogation of cancer-immune interactions. Given the well characterized role of immune surveillance in shaping the evolution of metastases^15,17–19^, this is expected to preclude discovery of important regulators of metastasis whose mechanism involves modulation of the immune response. Furthermore, they required post-hoc expansion of subclones identified in metastasis assays prior to characterization, limiting the fidelity of the captured cell states to the metastatic phenotypes observed.

To address these challenges, we generated subclones of mouse primary pancreatic ductal adenocarcinoma (PDAC) by capturing single cells and expanding them to produce monoclonal cell lines, each barcoded with a unique DNA sequence. We tested the relative metastatic potential of these subclones in competition assays in syngeneic immunocompetent mouse liver and peritoneum. These assays focused on the latter stages of the metastasis cascade, encompassing extravasating at the distant site, surviving the initial interactions with the metastatic microenvironment including the resident immune cells, and proliferating to form a macrometastasis. These latter steps of the metastatic cascade are the rate limiting steps^20^ and therefore the key to metastatic potential. Moreover, they are relevant to the adjuvant therapy setting when the primary tumor has been surgically removed and potential metastatic founders already distributed in the body. Our assays identified both metastasis-high and metastasis-low subclones, suggesting that we successfully captured primary tumor heterogeneity. We then performed ATAC-seq and RNA-seq on each subclone to respectively characterize its chromatin and transcriptomic states. This multi-omic characterization allowed us to connect the pre-metastatic state of the subclones to their ultimate metastatic performance.

We identified a set of metastasis-high (n = 207) and metastasis-low (n = 182) genes forming a metastatic potential axis orthogonal to the normal-to-PDAC and classical-to-basal axes. Metastasis-high genes were enriched for IL-1 pathway genes whereas metastasis-low genes were enriched for neuroendocrine, motility, and Wnt pathway genes. The majority of both the metastasis-high and -low genes are “pro-tumor”, meaning they are up-regulated in PDAC relative to normal pancreas, suggesting the met-high and -low states are separated predominantly by differential activation of shared pro-tumorigenic gene modules. In a targeted screen, we functionally validated the IL-1 pathway genes *Fos* and *Il23a* as well as other metastasis-high genes *Myo1b* and *Tmem40* as positive regulators of PDAC metastasis. Finally, to explore the relevance of our metastatic gene signature to human disease, we created the MetScore, a single sample score based on expression of metastasis-high and metastasis-low genes, and applied it to multiple publicly available human PDAC cohorts. We found that metastases have higher MetScores than primary tumors across cohorts and that higher primary tumor MetScores correlate with worse overall survival. Remarkably, MetScores were similarly higher in metastases than primary tumors amongst all other human carcinomas evaluated, suggesting the signature of metastatic potential we uncovered may be conserved across epithelial malignancies. This work establishes a strategy for using DNA barcodes to track metastatic performance of individual subclones in competition with others, and to link this performance to high-resolution multi-omic characterization of each subclone’s cell state. It further reveals important functional regulators of PDAC metastasis, and establishes a method for scoring human carcinomas based on metastatic potential.

## Results

### Isolation of primary PDAC subclones with high and low metastatic potential

To characterize the molecular determinants of metastatic potential within PDAC tumors, we obtained cells isolated from the primary tumors of two KPC mice^21–23^ (i.e., *Pdx1-Cre;LSL-Kras^G12D/+^;Trp53^R172H/+^*), which we denote here as KPC-1 and KPC-2. We barcoded each line by transposition of randomized DNA sequences^24,25^ (**Fig. 1A**). This strategy results in each cell receiving a unique combination of sequences that can act as a barcode for tracking it. From barcoded mixtures, we sorted single cells and expanded them to generate monoclonal lines. We obtained 10 barcoded subclones from KPC-1 and 7 from KPC-2 (**Fig. 1B, top**). We characterized the barcode in each subclone using high-throughput sequencing and cryopreserved several vials to enable downstream characterization and experimentation. In summary, we established 17 subclones from primary mouse KPC tumors, each with a unique and easily identifiable DNA barcode.

**Figure 1.**
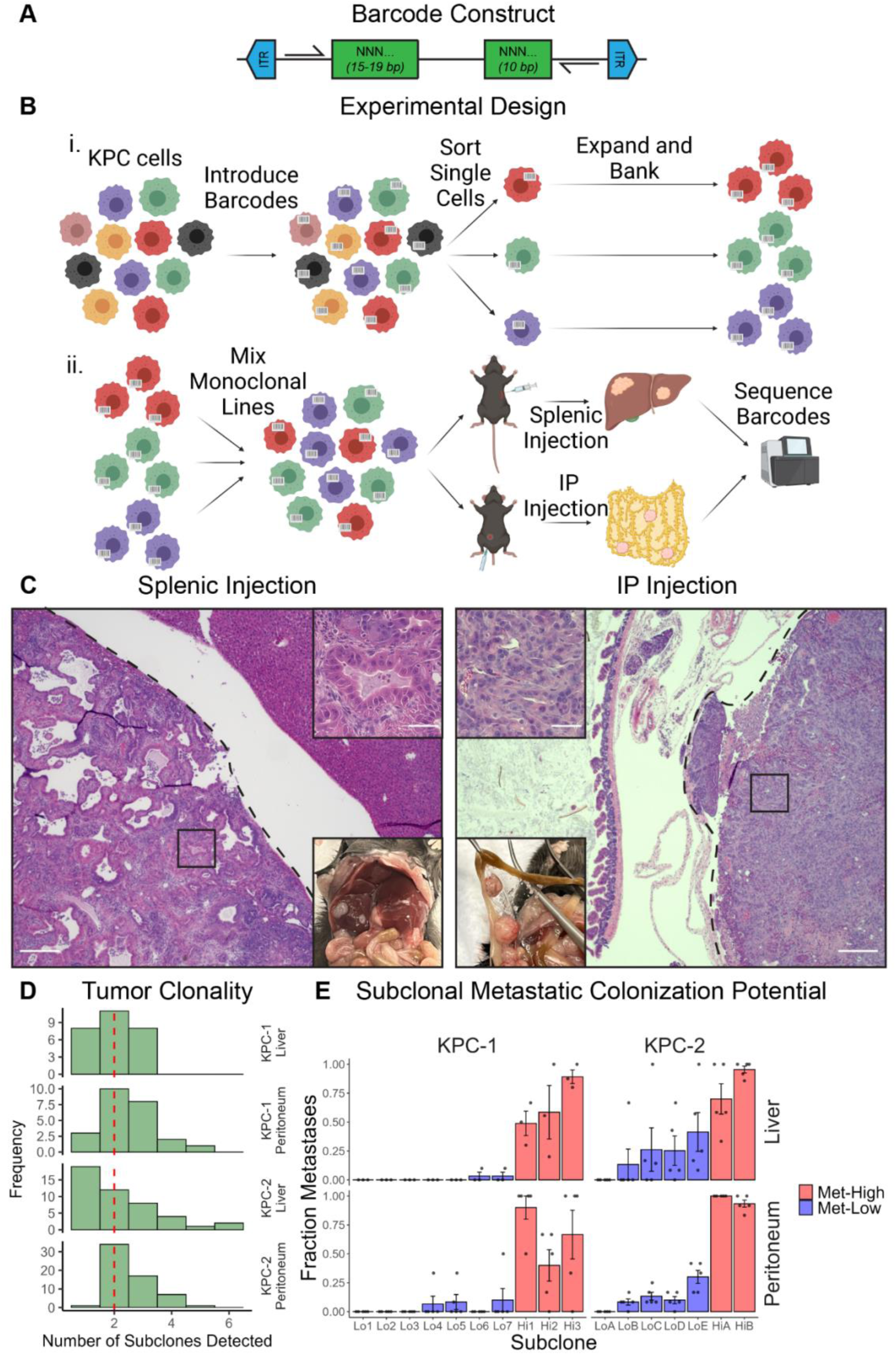
Isolation of primary PDAC subclones with high and low metastatic potential. (**A**) Schematic depicting barcode construct. (**B**) Schematic depicting process for producing barcoded monoclonal KPC lines and experimental design for *in vivo* metastasis competition assays. (**C**) Representative photographs and light micrographs of H&E stained FFPE sections. Scale bars indicate 500 µm on low magnification images and 50 µm on high magnification insets. (**D**) Histograms depicting distributions of tumor clonalities (i.e., the number of unique subclones detected in a given tumor) across the four experiments as denoted by the labels on the right-hand side. (**E**) Fractions of metastases in which each subclone was observed in each of the four experiments as denoted by the labels on the top and right-hand side. Individual mice represented by points and means across mice represented by bars with error bars representing S.E.M. (**D-E**) Sample sizes: KPC-1 liver, n = 27 mets across 3 mice; KPC-1 peritoneum, n = 24 mets across 5 mice; KPC-2 liver, n = 46 mets across 5 mice; KPC-2 peritoneum, n = 60 mets across 5 mice).

We next characterized the range of metastatic potentials of the isolated subclones. To do so, we introduced equal mixtures of all KPC-1 or KPC-2 subclones into immunocompetent syngeneic C57BL/6 mice via splenic and intraperitoneal routes, a standard approach for modeling metastasis to the liver^22,23^ and peritoneum^26^, respectively (**Figs. 1B,C**). After a 4-week incubation period, we harvested the resulting metastatic tumors and sequenced the barcode amplicon in each to identify its source subclone(s). The results showed that 80% of the sampled metastases were polyclonal, with a plurality being biclonal (**Fig. 1D)**. Therefore, we used the fraction of metastases in which a subclone is present as a measure of its metastatic potential. We did not use abundance in metastases as a measure to minimize confounding by growth potential after metastatic colonization. Metastatic potential showed a wide range in KPC-1 and KPC-2 subclones: some were present in nearly all metastases and others in nearly none (**Fig. 1E**). We denote subclones present in more than 50% of all analyzed metastases as ‘metastasis-high’ and the rest ‘metastasis-low’ subclones. Importantly, the subclones behaved consistently in both anatomic sites. This behavior indicates that the metastasis-high subclones captured here have a general advantage at forming metastases. Therefore, we have successfully deconvolved heterogeneous initial tumors into subclones with high or low metastatic potential.

The advantage of metastasis-high subclones can be based on their ability to complete the metastatic cascade. Alternatively, it can be based on a proliferative advantage for metastasis-high subclones. We assessed difference in proliferation in two ways. First, we performed *in vitro* proliferation assays (**Supp. Fig. 1A**). For both KPC-1 and KPC-2, the resulting growth curves were intermingled and did not exhibit separation between metastasis-high and metastasis-low subclones. Second, we performed *in vitro* competition assays mimicking our *in vivo* injections. We passaged equal mixtures of cells for the same length of time as *in vivo* growth and evaluated subclone dominance following the final passage by sequencing the barcode amplicon. Dominant subclones were strikingly different in the *in vitro* context compared to *in vivo* for both KPC-1 and KPC-2 (**Supp. Fig. 1B**), which is inconsistent with a growth advantage accounting for the observed *in vivo* metastatic potentials.

Another possible confounder for metastatic potential is that KPC tumors are heterogeneous and can include cells without *Kras* and *Trp53* mutations, which would likely present as metastasis-low in our assays. To test whether all isolated subclones are fully transformed, we genotyped all monoclonal lines and confirmed that they had all undergone recombination of the *LSL-Kras^G12D/+^* and *LSL-Trp53^R172H/+^*alleles (**Supp. Fig. 1C**). All monoclonal lines had also undergone loss of the wild-type *Trp53* locus (i.e., loss of heterozygosity [LOH]) with the exception of KPC-2_Lo1, a metastasis-low line. Because LOH at the *Trp53* locus is a pre-requisite for malignant transformation in the KPC mouse model^27^, we excluded KPC-2_Lo1 from further analyses to limit our cohort to only fully transformed subclones. Collectively, these results confirm that our metastasis-high PDAC subclones have an advantage in completing the metastatic cascade compared to the metastasis-low.

### The metastatic potential axis is orthogonal to the normal-to-PDAC and classical-to-basal axes

To understand the genesis of metastasis-high and metastasis-low subclones, we sought to place them in the natural history of PDAC development. We mapped the genome-wide chromatin accessibility landscape of the five metastasis-high and eleven metastasis-low subclones using ATAC-seq. We also obtained ATAC-seq data from epithelial cells derived from normal pancreata, pre-neoplasia from KC mice (i.e., *Pdx1-Cre;LSL-Kras^G12D/+^*), pancreatitis, pre-neoplasia with pancreatitis, and primary PDAC from KPC mice (KPC-0) that Alonso-Curbelo and colleagues^28^ generated in the same mouse strain as ours. To control for batch effects, we processed all the data (i.e., normal, pre-neoplasia, pancreatitis, pre-neoplasia+pancreatitis, primary PDAC, metastasis-low PDAC subclones, and metastasis-high PDAC subclones) using the same pipeline and then performed a batch correction on the KPC-1 and KPC-2 samples against KPC-0 using ComBat^29^. Our analysis strategy included alignment to the mouse genome, filtering of low quality, mitochondrial, and duplicate reads, and peak calling. We detected between 114,192 and 238,205 peaks per sample and all samples displayed greater than 10-fold enrichment of transcription start sites within their associated peaks, an important quality metric (**Supp. Table 3**). We then generated a consensus peak set common to all samples comprising 323,924 peaks by merging overlapping peaks from the individual peak sets. We scored each sample for all peak locations, batch corrected the KPC-1 and KPC-2 samples against KPC-0, and performed principal component analysis (PCA). Consistent with previous description, the results showed that normal, pancreatitis, pre-neoplasia, pancreatitis+neoplasia, and PDAC states were arranged sequentially along the first principal component (**Fig. 2A**), forming a normal-to-PDAC trajectory.

**Figure 2.**
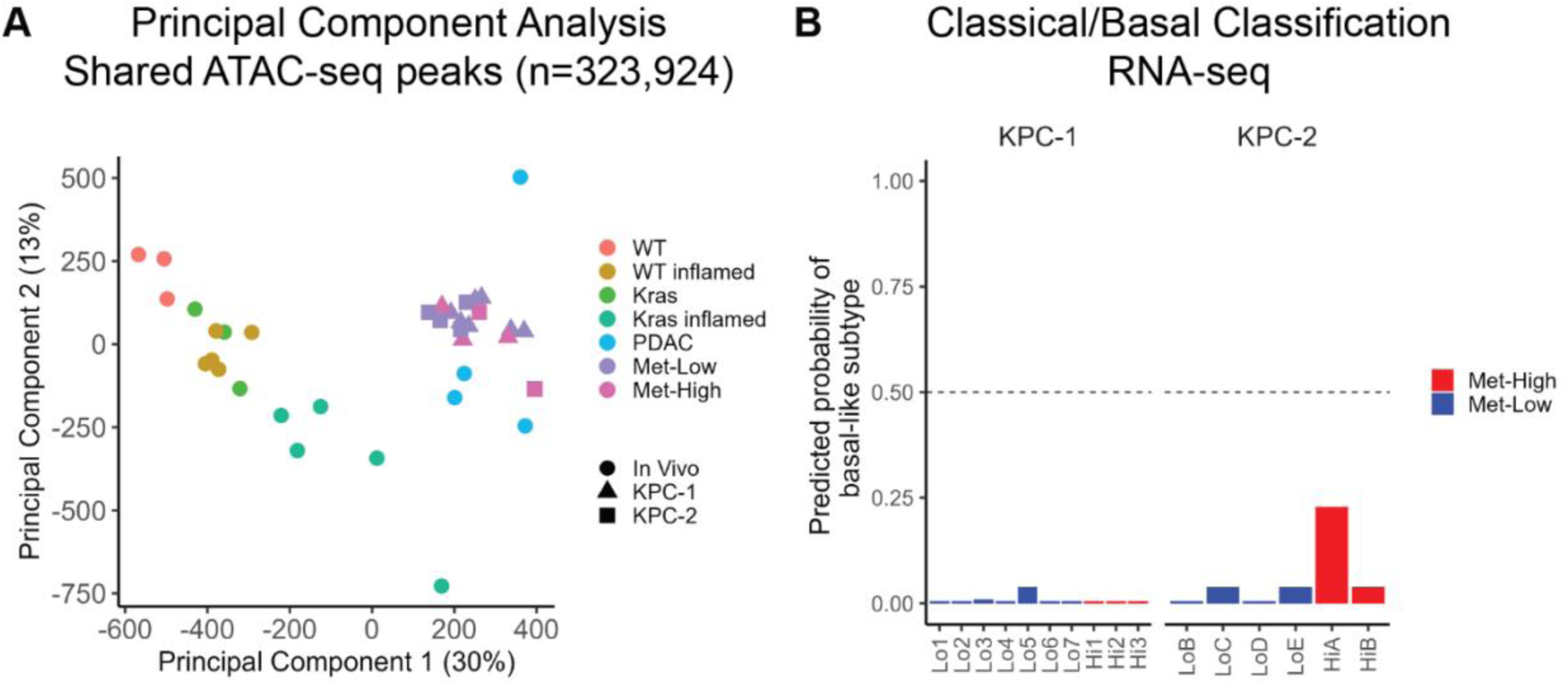
The metastatic potential axis is orthogonal to the normal-to-PDAC and classical-to-basal axes. (**A**) Principal component analysis of scaled normalized accessibility of a consensus ATAC-seq peak set. (**B**) Probabilities of basal-like molecular subtype occupancy for the metastasis-high and metastasis-low subclones based on application of the PurIST classifier to bulk RNA-seq data. A value less than 0.5 indicates likely classical state occupancy.

When we analyzed the position of our captured subclones in this normal-to-PDAC trajectory, we observed that metastasis-high subclones occupy the same overall state as metastasis-low subclones (**Fig. 2A**). The metastasis-high and metastasis-low subclones were intermingled without any separation between them along the principal components. To evaluate whether batch correction may have introduced bias masking global differences between metastasis-high and metastasis-low subclones, we performed PCA utilizing the single batch of data that contained normal, pancreatitis, pre-neoplasia, pancreatitis+pre-neoplasia, and primary PDAC samples only. We then transferred the loadings from the first two principal components to compare the metastasis-high and metastasis-low subclones in the context of these PCA axes (**Supp. Fig. 2**). Our subclones formed two clusters, largely concordant with their parental line, suggesting either biological or batch effect between KPC-1 and KPC-2. Still, this analysis did not separate metastasis-high and -low subclones, supporting our conclusion that metastasis-high and -low subclones occupy the same position along the normal-to-PDAC axis.

Landmark studies evaluating the transcriptomes of human primary PDAC tumors^30–32^ have identified recurrent molecular subtypes termed classical and basal, which have distinct molecular characteristics and clinical behavior. Subsequent single cell sequencing studies have deepened our understanding of this classification by demonstrating that within a single tumor, PDAC cells can display a spectrum of states existing along a classical to basal axis, with the fraction of each subtype varying between tumors^33^. To identify the molecular subtype of our captured subclones, we analyzed the transcriptomes of the five metastasis-high and 11 metastasis-low subclones using RNA-seq. We then classified each subclone as classical or basal utilizing a molecular classifier called PurIST^34^ that generates a single sample score representing the probability of basal-state occupancy from bulk transcriptomic data. PurlST categorized all metastasis-high and metastasis-low subclones as classical (**Fig. 2B**). Together, these results put forth a model wherein highly and poorly metastatic subclones arise from distinct branches during PDAC tumorigenesis that diverge along a metastatic potential axis orthogonal to the normal-to-PDAC and classical-to-basal axes.

### Identifying gene signatures for metastasis-high and metastasis-low states by integrating chromatin accessibility and gene expression

To characterize the chromatin features that define this metastatic potential axis, we identified ATAC-seq “peaks” (i.e., open chromatin regions) with differential accessibility between metastasis-high and metastasis-low subclones. To accomplish this, we generated a consensus peak set considering only the metastasis-high and metastasis-low samples, quantified the number of insertions at each peak location for each sample, and normalized the peak counts by the total reads aligning to peak regions for each sample. Principal component analysis demonstrated that the metastasis-high and -low samples capture the expected heterogeneity of different subclones within a primary tumor (**Fig. 3A**). We then applied a generalized linear model using DESeq2^35^ with the normalized peak insertion counts as the response variable, the metastatic potential (i.e., high vs. low) as a fixed effect, and parental group (i.e., KPC-1 vs. KPC-2) as a random effect. Out of the 176,964 total shared peaks, we identified 2,725 peaks (1.5%; **Supp. Table 4**) with increased accessibility in metastasis-high subclones and 5,720 peaks (3.2%; **Supp. Table 4**) with increased accessibility in metastasis-low subclones (**Figs. 3B-D**) using an FDR cutoff of 0.05. The majority of the differential peaks fall in distal intergenic regions (40.9%) or introns (36.6%), and thus are likely to represent enhancers (**Fig. 3E**). Indeed, 76.3% of the differentially accessible peaks overlap candidate regulatory elements (REs) identified by the ENCODE project and 62.7% of all peak-RE overlaps involve candidate enhancers (**Fig. 3F**). The plurality of the remaining peaks fall in promoter regions (16.9%; **Fig. 3E**) and a small number fall in exons (4.0%). The observation that most of the identified differential peaks are located in putative enhancer regions aligns with the prevailing model of gene regulation, which posits that enhancers play a critical role in cell type-specific gene expression^36^; in contrast, promoters are generally more consistently accessible across different cell types, contributing to a uniform regulation of gene expression. In summary, we identified 8,445 peaks in our ATAC-seq data whose accessibility differed between metastasis-high and metastasis-low PDAC subclones.

**Figure 3.**
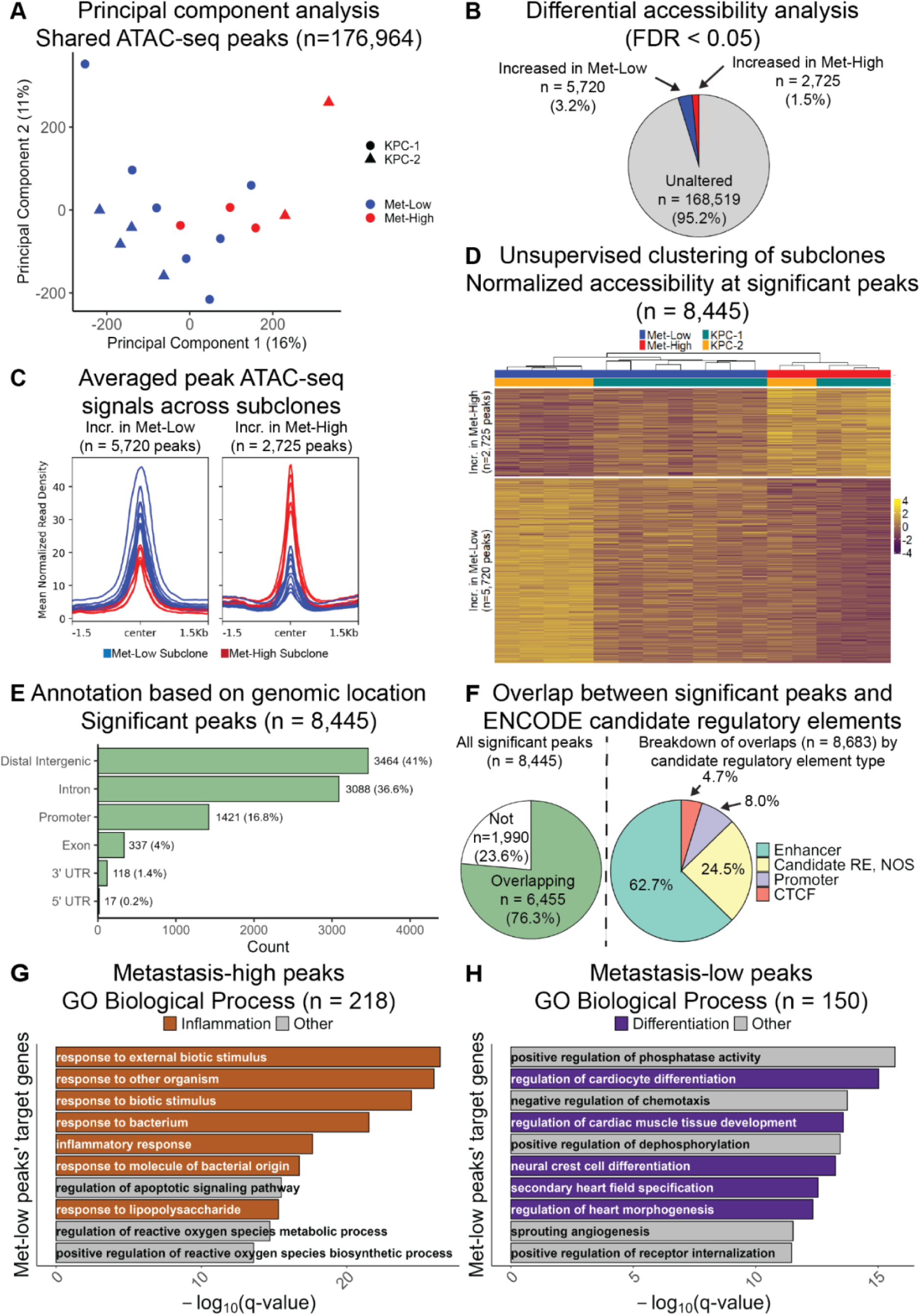
Identifying differentially accessible open chromatin regions specific to metastasis-high and metastasis-low PDAC subclones. (**A**) Principal component analysis of normalized accessibility of a consensus peak set. (**B**) Pie chart illustrating the fraction of total peaks in the consensus peak set found to have significantly different accessibility between metastasis-high and metastasis-low subclones when controlling for parental group status using the generalized linear model feature of DESeq2. An FDR cutoff of 0.05 was used. (**C**) Averaged ATAC-seq signals for peaks with increased accessibility in metastasis-low subclones (left) and increased accessibility in metastasis-high subclones (right). Each line represents a subclone colored based on its metastatic potential with red indicating high and blue indicating low (**D**) Heatmap depicting the normalized accessibility for each differential peak (row) for each subclone (column). Subclones were clustered based on Pearson correlation (**E**) Bar plot depicting the breakdown of genomic locations for the identified significant differentially accessible peaks with respect to gene annotations. Promoter here is defined as the region up to 3 kb upstream of the transcription start site. (**F**) Pie charts depicting (left) the fraction of significant differentially accessible peaks overlapping candidate regulatory elements (REs) identified by the ENCODE project and (right) the breakdown for the overlaps with respect to the candidate RE type. (**G-H**) Bar plots demonstrating the most significant GO Biological Process pathways when applying Genomic Regions Enrichment of Annotations Tool (GREAT) to significant peaks with increased accessibility in metastasis-high (**G**) or metastasis-low (**H**) subclones. Default settings for GREAT were used. Pathways ranked in order of decreasing significance based on binomial FDR *q*-value. Pathways meeting a threshold of FDR < 0.05 for both binomial and hypergeometric tests were considered significant.

The epigenetic comparison above identified chromatin features which segregate with observed differences in metastatic potential. Since metastatic potential is mitotically heritable, we expect its underlying features to be among these chromatin features. To link the differentially accessible peaks to biological pathways that may be important for metastasis, we evaluated whether the metastasis-high and metastasis-low peaks are enriched in regulatory regions of genes in Gene Ontology^37,38^ (GO) Biological Process pathways using GREAT^39^ (Genomic Regions Enrichment of Annotations Tool). There were 218 GO pathways enriched amongst the metastasis-high peaks’ target genes (**Fig. 3G**; **Supp. Table 5**), with seven out of the top 10 being related to inflammation (e.g., “inflammatory response”), and there were 150 GO pathways enriched amongst the metastasis-low peaks’ target genes (**Fig. 3H**; **Supp. Table 5**), with five out of the top ten being related to tissue or organ development or differentiation (e.g., “regulation of cardiocyte differentiation”), considering only pathways passing an FDR cutoff of 0.05 for both binomial and hypergeometric tests. While this analysis likely points to some pathways that are important for metastasis, given the large number and diversity of pathways identified, it is unlikely that all of the identified pathways are functionally related to metastatic potential.

To narrow down this list to a high confidence set of genes more likely to be functionally related to metastasis, we reasoned that in our assay to measure metastatic potential (**Fig. 1B**), successful cells must realize their advantage in completing the latter half of the metastatic cascade immediately after injection; therefore, these cells are likely to be expressing the genes that confer them this advantage at the time of injection. To identify which differentially accessible open chromatin regions were also associated with gene expression differences between metastasis-high and metastasis-low subclones, we used the RNA-seq data generated from the five metastasis-high and eleven metastasis-low subclones. Again, we utilized the generalized linear model functionality of DESeq2, this time to isolate differentially expressed genes between metastasis-high and metastasis-low subclones while controlling for parental group status, identifying a total of 932 significant genes, 498 with increased expression in metastasis-high subclones and 433 with increased expression in metastasis-low subclones (**Fig. 4A**; **Supp. Table 6**) using an FDR cutoff of 0.05. Next, we assigned ATAC-seq peaks to their nearest gene and observed correlation of differentially accessible peaks with differential gene expression (**Fig. 4B**). 84% of differentially accessible peaks were assigned to genes with unchanged expression after filtering out 124 peaks assigned to predicted genes lacking an Ensembl ID and 2,640 peaks linked to genes that were lowly expressed in both metastasis-high and metastasis-low subclones. This result is expected as some peaks may not regulate any genes, their target gene may be in a poised state, or their target may not be the nearest gene. For differentially accessible peaks linked to differentially expressed genes, the genes tended to follow the expected pattern, with 97% of significant genes linked to peaks with increased accessibility in metastasis-high subclones having increased expression in metastasis-high subclones, and 91% of significant genes linked to peaks with increased accessibility in metastasis-low subclones having increased expression in metastasis-low subclones. All peaks assigned to the top 50 genes with significantly higher expression in metastasis-low subclones had greater accessibility in metastasis-low subclones and a substantial majority (i.e., 79%) of peaks assigned to the top 50 genes with significantly higher expression in metastasis-high subclones had greater accessibility in metastasis-high subclones (**Fig. 4C**). We define the genes with differential expression concordant with the differential accessibility of at least one assigned peak to be “metastasis-high genes” (n = 207) or “metastasis-low genes” (n = 182). To highlight two representative examples, multiple peaks in the promoter region and first intron of *Il18r1*, a metastasis-high gene, displayed increased accessibility in concert with increased expression of *Il18r1* in metastasis-high subclones (**Fig. 4D**), and conversely multiple peaks in the proximal and distal intergenic regions surrounding *Mnx1*, a metastasis-low gene, as well as its promoter region displayed increased accessibility in concert with increased expression of *Mnx1* in metastasis-low subclones (**Fig. 4E**).

**Figure 4.**
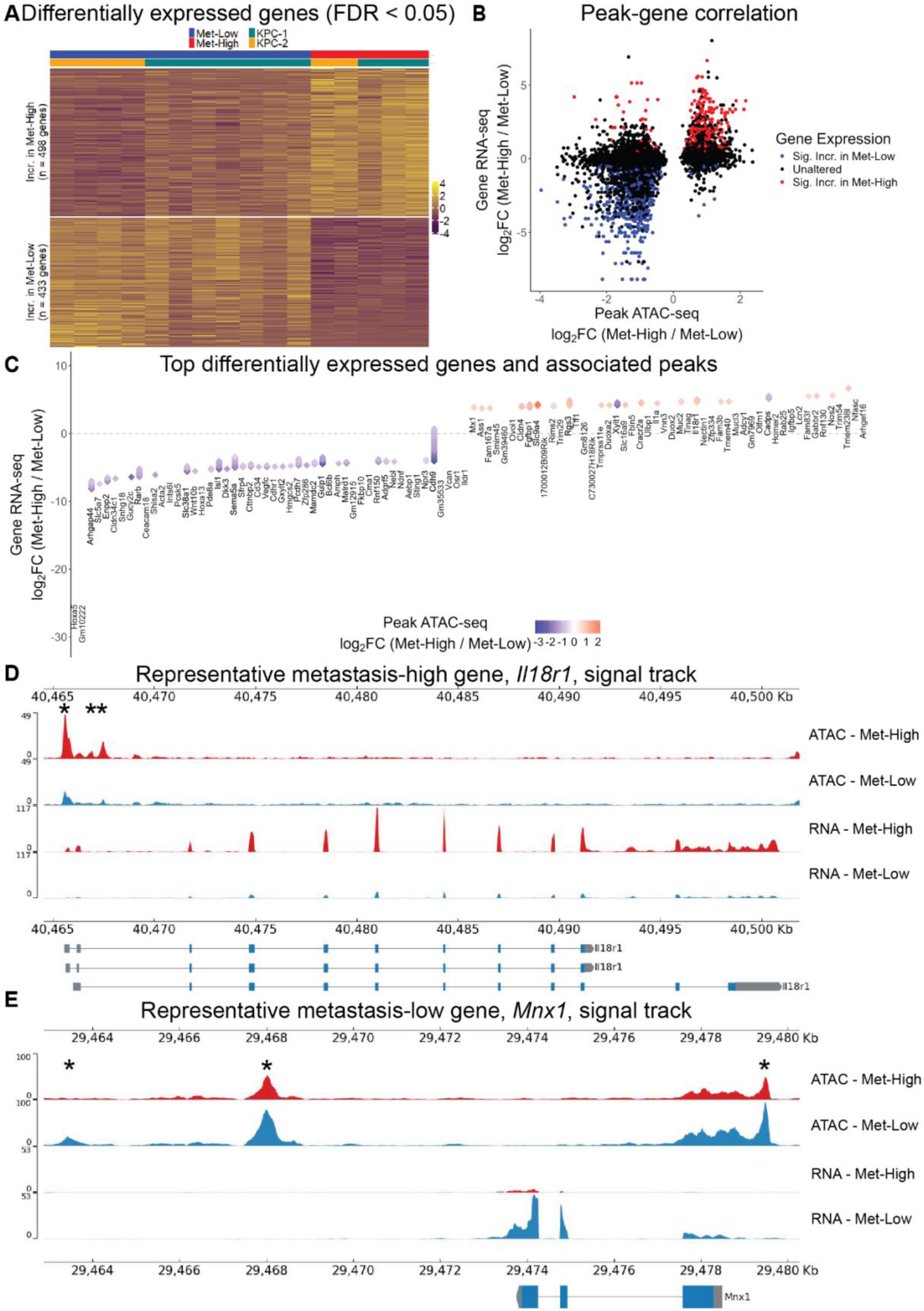
Isolating metastasis-high and metastasis-low defining genes by integrating chromatin accessibility and gene expression. (**A**) Heatmap depicting normalized expression for each significantly differentially expressed gene (row) for each subclone (column). An FDR cutoff of 0.05 was used. (**B**) Scatterplot depicting, for each significantly differentially accessible peak (see **Fig. 3**), that peak’s differential accessibility between metastasis-high and metastasis-low subclones on the X axis and the nearest gene’s differential expression between metastasis-high and metastasis-low subclones on the Y axis. The color of the peak indicates whether the nearest gene is significantly differentially expressed. (**C**) Diamond plot depicting the top 50 most downregulated and top 50 most upregulated genes in metastasis-high subclones relative to metastasis-low. The position of the gene label on the Y axis indicates that gene’s differential expression. The genes are arranged by rank from most downregulated in metastasis-high on the left to most upregulated in metastasis-high on the right. Above each gene label are arranged diamonds representing significantly differentially accessible peaks for which the noted gene is the closest gene. The color of the diamond indicates that peak’s normalized differential accessibility between metastasis-high and metastasis-low subclones. (**D-E**) Signal tracks depicting chromatin accessibility and gene expression of genomic regions containing *Il18r1* or *Mnx1*, representative metastasis-high and metastasis-low genes, respectively, in the metastasis-high and metastasis-low subclones. Asterisks indicate peaks identified to be differentially accessible between metastasis-high and metastasis-low subclones.

In summary, we have identified a bi-directional gene signature associated with the metastatic potential of PDAC subclones: metastasis-high genes show higher relative expression in subclones with a high metastatic potential while metastasis-low genes are relatively increased in subclones with a lower metastatic potential. Because each of these genes is also associated with chromatin regions showing a concordant degree of openness, they are more likely to be under stable epigenetic regulation required for the mitotic transmission of metastatic potential.

### IL-1 pathway genes are enriched amongst metastasis-high genes and neuroendocrine, motility, and Wnt pathway genes are enriched amongst metastasis-low genes

The metastasis-high and metastasis-low genes represent our most confident sets of genes that can contribute to metastatic potential. To gain insights into the biological functions enabled by these genes, we applied gene set enrichment analysis to the metastasis-high and metastasis-low sets using the GO^37,38^ Biological Process and Kyoto Encyclopedia of Genes and Genomes^40^ (KEGG) Pathway databases. The top three out of the five total KEGG gene sets significantly enriched within the metastasis-high genes using an FDR cutoff of 0.05 are related to infection or inflammation (i.e., “TNF signaling pathway”, “Pertussis”, and “Leishmaniasis”; **Fig. 5A; Supp. Table 8**). No GO terms were enriched within the metastasis-high genes using an FDR cutoff of 0.05. Further examination indicated that these GO terms coalesce on the IL-1 signaling pathway: Nearly all (9/11) of the genes present in these infection/inflammation gene sets (**Fig. 5B**) fall within this pathway, including an IL-1 family cytokine (*Il1a*)^41^ and cytokine receptor (*Il18r1*)^41^, machinery for converting extrinsic IL-1 signals into NF-κB activity (*Irak1*, *Tab3*)^42^, and well-established NF-κB transcriptional targets (*Nos2*, *Il1a*, *C3*, *Cxcl5*, *Fos*, *Xiap*)^43–48^. We define an “IL-1 Pathway” module as the set comprising these nine genes (**Fig. 5E**).

**Figure 5.**
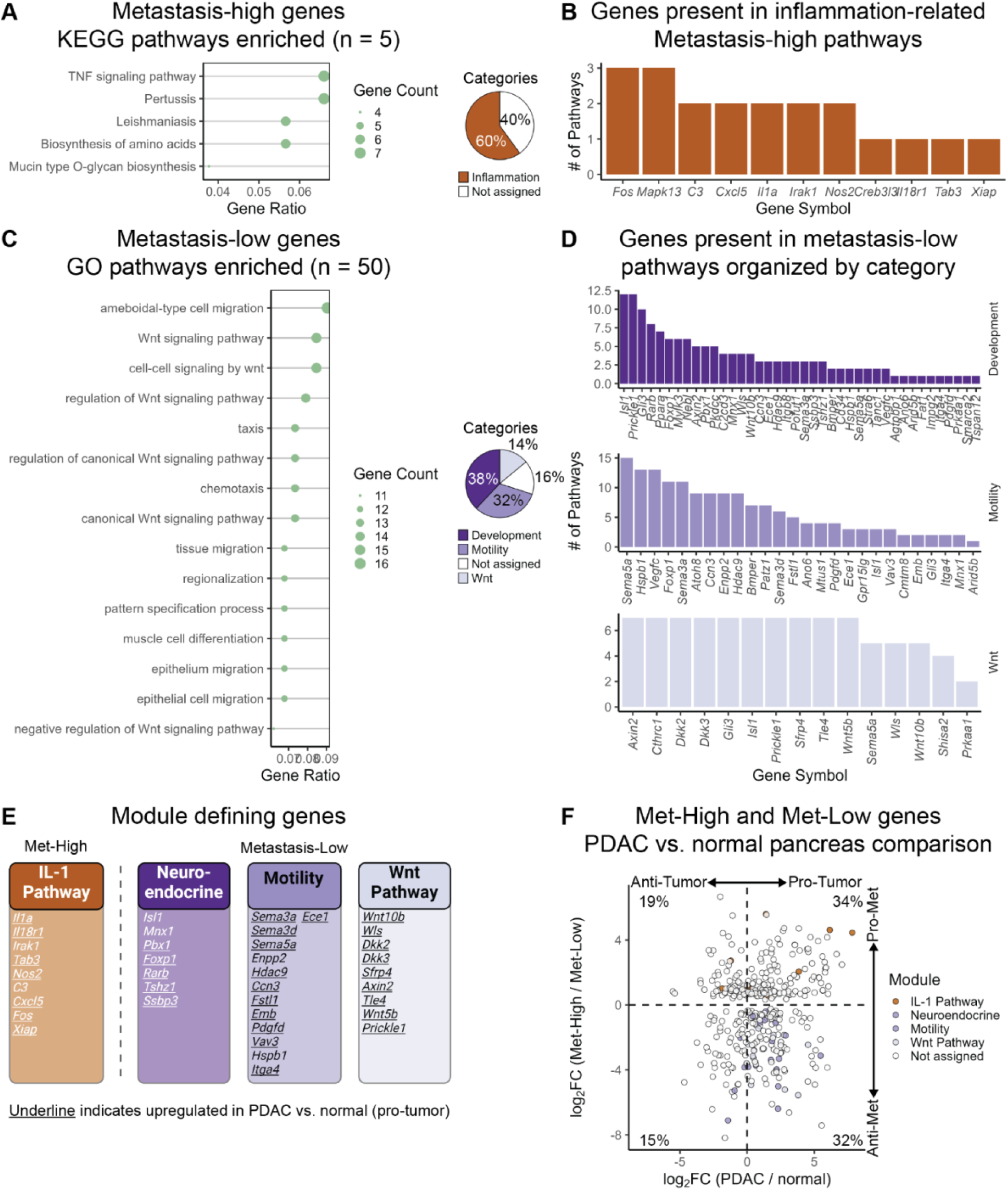
IL-1 pathway genes are enriched amongst metastasis-high genes and neuroendocrine, motility, and Wnt pathway genes are enriched amongst metastasis-low genes. (**A**) The left panel is a dotplot depicting gene ratio and gene count (i.e., the number of metastasis-high genes present in the pathway in question), for all significantly enriched KEGG pathways amongst the metastasis-high genes using an FDR cutoff of 0.05. The right panel is a pie chart depicting the fraction of enriched pathways manually curated as being related to inflammation (**B**) Bar plot depicting, for all of the genes found within any inflammation-related pathway enriched amongst metastasis-high subclones, the number of inflammation-related pathways in which it is found. (**C**) The left panel is a dotplot depicting gene ratio and gene count for all significantly enriched GO pathways amongst the metastasis-low genes using an FDR cutoff of 0.05. The right panel is a pie chart depicting the fractions of enriched pathways manually curated as being related to development, motility, or Wnt. (**D**) Bar plot depicting, for all of the genes found within any pathway within the denoted manually curated groups of pathways (i.e., development, motility, or Wnt), the total number of pathways in that manually curated group of pathways in which that gene is found. (**E**) Schematic depicting genes defining each of the four metastasis-high and metastasis-low specific gene modules. (**F**) Scatterplot depicting each gene in the metastasis-high and metastasis-low gene sets. Position on the X axis indicates log_2_ fold change between primary PDAC and normal pancreas samples. Position on the Y axis depicts log_2_ fold change between metastasis-high and metastasis-low subclones. Color indicates that gene’s membership in one of the gene modules defined in **E**.

Amongst metastasis-low genes, there were 50 GO pathways significantly enriched using an FDR cutoff of 0.05 (**Fig. 5C; Supp. Table 8**). These pathways were divided by manual curation into three classes: development (19/50; e.g., “muscle cell differentiation”); motility (16/50; e.g., “ameboidal-type cell migration”); and Wnt (7/50; e.g., “Wnt signaling pathway”). The remaining eight pathways could not be confidently assigned into one of these groups. No KEGG terms were enriched within the metastasis-low genes using an FDR cutoff of 0.05. Review of metastasis-low genes present in the development gene sets identified multiple genes required for proper islet development including *Isl1*^49^, *Mnx1*^50^, *Pbx1*^51^, *Foxp1*^52^*, Rarb*^53^, *Tshz1*^54^, and *Ssbp3*^55^ (**Fig. 5D**), which we collectively define as the “Neuroendocrine” module (**Fig. 5E**). This observation is consistent with the “developmental constraint” model of cancer evolution^56^, which posits that cancer cells evolve by accessing gene programs specific to developmental lineages adjacent to the cell of origin, which in this case is controversial but is most likely either the mature pancreatic acinar cell or mature pancreatic ductal cell^57^. Furthermore, this observation is consistent with a prior observation of subpopulations of human PDAC cells in a neuroendocrine-like differentiation state^58^.

Metastasis-low genes found in motility gene sets include the semaphorins *Sema3a*, *Sema3d*, and *Sema5a*, which function under normal physiology as axon guidance cues^59^ (**Fig. 5D**). Semaphorins are generally believed to have a tumor-promoting role in PDAC development. *SEMA3A* and *SEMA3E* are recurrently amplified in human PDAC^60^. Experiments performed in mouse models suggest that semaphorins promote metastasis both through cancer-cell intrinsic mechanisms such as stimulating migration^61,62^ and cancer-cell extrinsic mechanisms such as polarizing macrophages towards an M2 phenotype^63^ and recruiting neurons^64^. The metastasis-low genes found in motility gene sets also include a number of genes that have been shown to stimulate motility of cancer cells under certain circumstances including *Enpp2*^65^, *Hdac9*^66^, *Ccn3*^67^, *Fstl1*^68^, *Emb*^69^*, Pdgfd*^70^, *Vav3*^71^, *Hspb1*^72^, *Itga4*^73^, and *Ece1*^74^. We define a “Motility” module as the set comprising these ten genes as well as the three above mentioned semaphorin genes (**Fig. 5E**). While we did not initially expect to observe motility-related genes enriched in the metastasis-low subclones given the general assumption that a motile state is advantageous for a cell to complete the metastatic cascade, as we will show below, these genes are still upregulated in the metastasis-high subclones compared to normal pancreatic epithelium. Therefore, one possible explanation is that metastasis requires a moderate level of increased motility but that extreme dysregulation is inhibitory. Another possible explanation is that motility may be important for egress from the primary tumor but dispensable for the latter stages of the metastatic cascade that are being captured by our assays. Yet another possible explanation is that certain pro-motility genes, which may be metastasis enabling, are co-regulated with a broader set of genes that are anti-metastatic.

Metastasis-low genes found in Wnt pathway gene sets include the canonical Wnt ligand *Wnt10b*^75^ (**Fig. 5D**) as well as *Wls*, whose gene product WLS facilitates the secretion of canonical Wnt ligands. However, there are also a number of genes encoding canonical Wnt pathway antagonists amongst the metastasis-low genes (*Dkk2*, *Dkk3*, *Sfrp4*, *Axin2*, and *Tle4*)^75^. In addition, a gene encoding a non-canonical Wnt ligand, *Wnt5b*^75^, as well as an essential planar cell polarity gene, *Prickle1,* are found amongst the metastasis-low genes. We define a “Wnt” module as the gene set comprising these nine genes (**Fig. 5E**). This is challenging to interpret given the presence of both agonists and antagonists amongst the metastasis-low genes as well as both canonical Wnt and non-canonical Wnt/planar cell polarity genes but suggests differential dysregulation of the Wnt pathways between the metastasis-high and metastasis-low subclones.

Above, we identified genes and pathways with different expression levels in metastasis-high and metastasis-low PDAC subclones relative to each other. An important missing context of this analysis is that the directionality of gene expression changes relative to the baseline normal pancreas state is unclear. To address this, we used RNA-seq data from epithelial cells derived from normal pancreata and primary PDAC from KPC mice (KPC-0), again generated by Alonso-Curbelo and colleagues^28^ using the same mouse strain as ours. We performed differential gene expression analysis using the KPC-0 and normal pancreata samples, allowing us to identify each metastasis-high and metastasis-low gene as down-regulated in tumor (anti-tumor) or up-regulated in tumor (pro-tumor). Combining this information with the enrichment level of each gene in metastasis-high versus metastasis-low subclones allowed us to place each gene on both an anti-metastasis to pro-metastasis axis and an anti-tumor to pro-tumor axis (**Fig. 5F; Supp. Table 9**). The majority of both the metastasis-high (133/207 [64.3%]) and the metastasis-low (123/182 [67.6%]) genes were identified as being pro-tumor. Thus, most of the differences between the two states are related to genes being activated during primary tumorigenesis to a more or less pronounced degree in each state. The IL-1 Pathway module genes are largely (82%) pro-tumor (*Nos2*, *Il18r1*, *Il1a*, *Cxcl5*, *Fos, Tab3*, and *Xiap*) with the exception of *C3* and *Irak1*, which are anti-tumor. Similarly, 5/7 (71.4%) of the Neuroendocrine module genes, 11/13 (84.6%) of the Motility module genes including all three semaphorin genes, and all (9/9) Wnt Pathway module genes are pro-tumor. Taken together, these results suggest that a majority of metastatic potential is regulated by differential prioritization of pathways that are already important for tumorigenesis. They also identify candidate genes which may be positively contributing to initial tumorigenesis but dispensable or detrimental to metastatic potential.

### NF-κB and mesenchymal transcription factors regulate the metastasis-high state whereas CDX2 and HOXA13 regulate the metastasis-low state

Coordinated activation of multiple genes in each module associated with metastasis-high or metastasis-low subclones points to transcription factors (TFs) helping drive or maintain these states. To identify TFs associated with the metastasis-high and metastasis-low states, we used TF footprinting^76^, which searches for sites within ATAC-seq peaks that show a dip in openness signal. Such dips can be attributed to a TF binding to the DNA at that site and preventing the Tn5 transposase from inserting in that position, thereby creating a ‘footprint’ within the peak. We first merged aligned ATAC-seq reads from the five metastasis-high subclones and, separately, from the eleven metastasis-low subclones after downsampling the reads for each to match the least covered subclone in that group to ensure equal representation of each subclone. We then utilized the package TOBIAS^76^ to identify footprints and calculate the differential binding scores (DBSs) for 511 TF motifs in the JASPAR database^77^ whose TFs are expressed across the tested subclones. There were 209 (∼41%) motifs with significantly greater DBSs in metastasis-high subclones and 290 (∼57%) motifs with significantly greater DBSs in metastasis-low subclones (Bonferroni adjusted *p*-value < 0.05; **Fig. 6A; Supp. Table 10**). Validating these predicted sets, motifs corresponding to the four TFs involved in islet development previously identified as metastasis-low genes, i.e., *Isl1*, *Mnx1*, *Pbx1* and *Foxp1*, all had DBSs favoring the metastasis-low subclones (two-sided T test, adjusted *p*-value < 1E-16 for all four). Furthermore, 4 of 6 motifs assigned to the inflammation-related metastasis-high gene *Fos* had DBSs favoring the metastasis-high subclones (two-sided T test, adjusted *p*-value < 1E-16 for FOS motif 1, FOS::JUN motif 1, FOS::JUNB, FOS::JUND).

**Figure 6.**
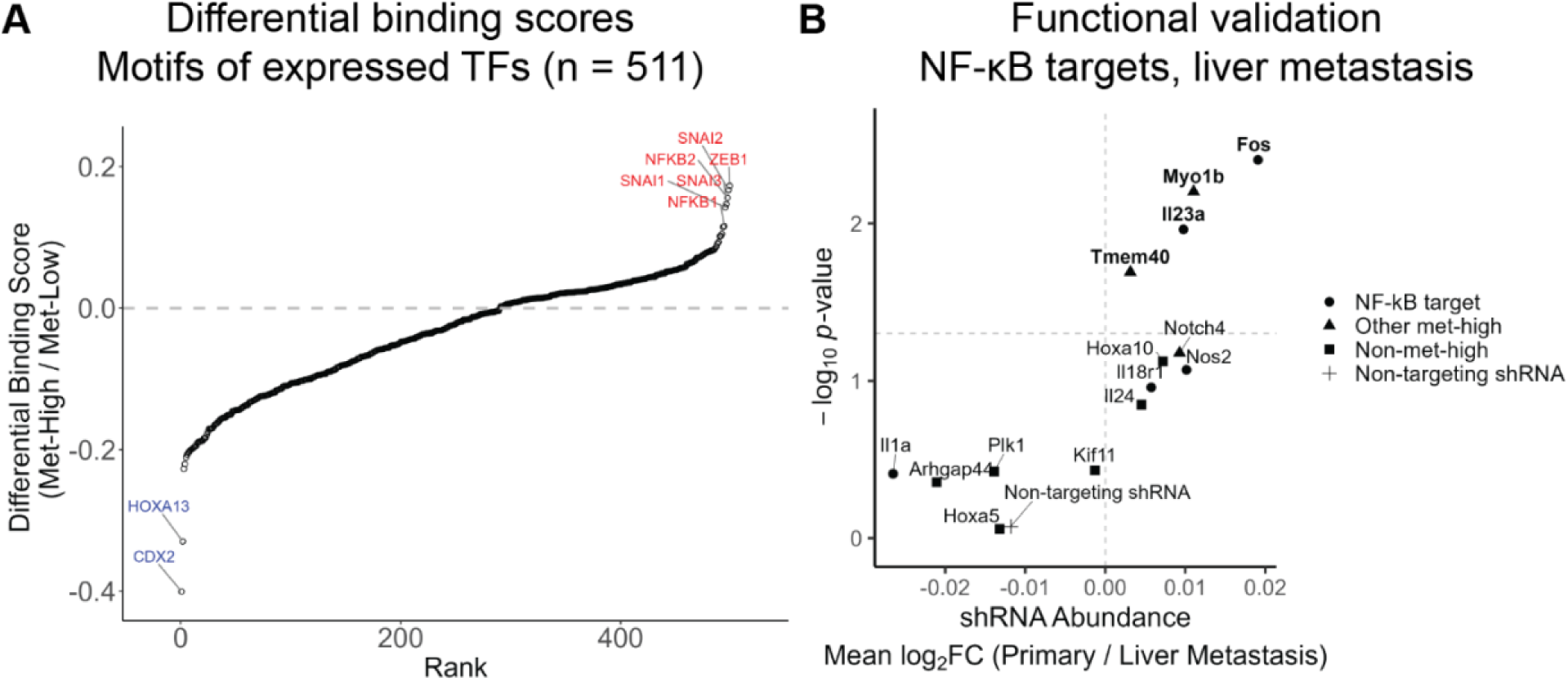
NF-κB and mesenchymal transcription factors regulate the metastasis-high state whereas CDX2 and HOXA13 regulate the metastasis-low state. (**A**) Rank-ordered plot of differential binding scores for significant transcription factor motifs using a Bonferroni adjusted *p*-value cutoff of 0.05. Only motifs in the JASPAR CORE vertebrates nonredundant set expressed in the KPC subclones were considered (n = 511) and only motifs with significant DBSs (n = 499) are being shown here. Motifs are arranged on the x-axis by rank according to their differential binding scores. The y-axis represents the differential binding score, with positive values indicating increased binding in metastasis-high subclones and negative values indicating increased binding in metastasis-low subclones. Motifs with the greatest effect sizes are highlighted. (**D**) Scatterplot depicting genes included in a targeted shRNA screen with position along the X axis representing mean *log*_2_ fold change in abundance amongst all shRNAs targeting each gene between primary tumor and liver conditions and position along the Y axis representing the −*log*_10_ weighted combined *p-*value generated using a linear model. Sample sizes: six primary tumors across six mice and 36 liver metastases across four mice.

The top motifs with differential binding scores favoring the metastasis-low subclones belong to CDX2 and HOXA13 (two-sided T test, adjusted *p-*value < 1E-16 for both), which are both involved in anterior-posterior patterning during development. CDX2, part of the ParaHox cluster, is expressed in gut epithelium caudal to the stomach beginning with hindgut invagination and continuing throughout development and into adulthood and functions to suppress the default state of forestomach endoderm and promote a caudal phenotype^78^. Hox family members are generally not expressed in gut endoderm but are expressed primarily in developing ectodermal and mesodermal tissues and are responsible for segmental specification of the musculoskeletal and nervous systems^78^. HOXA13 may be a rare exception to this rule as it has been shown to be expressed in the endoderm of the cloacal region of the developing chick^79^. Both *Cdx2* and *Hoxa13* genes have increased expression in metastasis-low subclones relative to metastasis-high subclones (log_2_ fold change [metastasis-high / metastasis-low] −1.771, Wald test adjusted *p*-value 6.5E-4 for *Cdx2*; log_2_ fold change −5.549, adjusted *p*-value 2.6E-3 for *Hoxa13*). These results are again consistent with the developmental constraint model of cancer evolution^56^ in which cancer cells evolve by co-opting gene modules belonging to developmental lineages adjacent to the cancer’s cell of origin, in this case navigating in the reverse direction along the developmental map from the mature acinar or ductal cell to the pancreatic progenitor cell to primitive gut endoderm. They also suggest that the transcriptional program activated by Hox and ParaHox transcription factors may blunt the metastatic potential of PDAC subclones.

The top motifs with differential binding scores favoring the metastasis-high subclones belong to the mesenchymal TFs ZEB1, SNAI2, SNAI3, and SNAI1 and NF-κB subunits NFKB2 and NFKB1 (two-sided T test, adjusted *p-*value < 1E-16 for all five). The Zeb and Snail family transcription factors promote epithelial-to-mesenchymal transition (EMT) in the context of carcinomas such as PDAC^80^. Maintenance of E-cadherin expression in the metastasis-high subclones compared to the metastasis-low (Wald test, adjusted *p*-value 0.23 for *Cdh1*) points to the metastasis-high subclones being in a hybrid-EMT state. This is consistent with a prior lineage tracing study demonstrating that PDAC subclones in hybrid-EMT states have greater metastatic potential than subclones in fully epithelial or fully mesenchymal states^12^. There are five NF-κB subunits^42^ including the two previously mentioned and all five have associated motifs with DBSs favoring the metastasis-high subclones (adjusted *p*-value < 1E-16 for REL, RELA, and RELB). This is consistent with our prior discovery that *Il1a*, *Il18r1* (an IL-1 family receptor), and multiple validated NF-κB transcriptional targets (*Nos2*, *Il1a*, *C3*, *Cxcl5*, *Fos*, and *Xiap*)^43–48^ have increased expression in the metastasis-high subclones in concordance with an open state of chromatin at their nearby cis-regulatory elements (**Figs. 4C-D & 5A-B**). Overall, these results identify NF-κB, Zeb, and Snail families as the most prominent among the transcriptional regulators of the metastasis-high subclones, pointing to inflammatory response and EMT as important modulators of metastatic potential.

To evaluate the functional impact of NF-κB pathway activity on metastatic potential, we performed a targeted shRNA screen that included NF-κB target genes (*Il1a*, *Il18r1*, *Nos2*, *Fos*, *Il23a*), other metastasis-high genes (*Tmem40*, *Myo1b*, *Notch4*), and a few non-metastasis-high genes from the mouse genome to serve as controls. We focused on liver metastasis since liver is the most common site of human PDAC metastasis and liver metastasis portends an especially poor prognosis^81^. We infected the KPC-1_Hi2 subclone with a lentiviral shRNA library containing three unique shRNAs targeting each gene as well as three control non-targeting shRNAs. We then used these cells to generate liver metastases by performing splenic injections. For our control condition, we generated primary tumors via orthotopic injection to separate general pro-tumor effects, such as proliferation and engraftment, from specific contributions to liver metastasis. Following an incubation period, we harvested the resulting tumors and sequenced the shRNA amplicon contained therein. The results show that shRNAs targeting *Fos* and *Il23a* were significantly depleted in liver metastases relative to primary tumors (linear model, weighted combined *p*-value 3.9E-3 for *Fos* and 0.011 for *Il23a*), indicating that these NF-κB target genes involved in the IL-1 pathway are novel functional enablers of metastasis and consistent with an overall positive effect of NF-κB pathway activity on metastatic potential. In addition, shRNAs targeting the metastasis-high genes *Myo1b*, which encodes an atypical myosin^82^, and *Tmem40*, which encodes a transmembrane protein with poorly characterized function^83^, were similarly significantly depleted in liver metastases relative to primary tumors (weighted combined *p-*value 6.3E-3 for *Myo1b* and 0.020 for *Tmem40*), validating these two additional genes as novel functional enablers of metastasis and, in combination with the results for *Fos* and *Il23a*, providing confidence that our approach is an effective strategy for nominating candidate functional enablers.

Taken together, the results so far paint a complex picture of factors associated with metastatic potential, which involves hundreds of genes and tens of transcription factors. However, a clear feature of the metastasis-high subclones is the activation of inflammation-related genes while metastasis-low subclones show activation of gene expression programs specific to adjacent development lineages, specifically pancreatic endocrine progenitor and primitive gut endoderm.

### Metastasis-high and metastasis-low genes define a metastasis signature in human carcinomas

We next assessed the relevance of our bi-directional metastatic gene signature in the KPC mouse model to human PDAC. We focused on two cohorts of human PDAC with gene expression profiles for primary and metastatic tumors available: the International Cancer Genome Consortium PACA-CA cohort, which includes RNA-seq from tumor biopsies harvested from patients with untreated locally advanced or metastatic PDAC enrolled in the COMPASS^84^ and PanGen^85^ trials, and a second cohort herein denoted as “PACA-US” that includes microarray data from primary and metastatic tumors harvested from deceased individuals with PDAC enrolled in the Nebraska Medical Center Rapid Autopsy Pancreatic Program and the Johns Hopkins Gastrointestinal Cancer Rapid Medical Donation Program as well as resected primary PDAC from living patients at the Johns Hopkins Medical Institutions, Northwestern Memorial Hospital, NorthShore Hospital and UNC hospitals^31^. We reasoned that if primary tumor subclones with elevated expression of metastasis-high genes and diminished expression of metastasis-low genes indeed have increased odds of seeding distant metastases, the gene signature is likely to persist, at least partially, in the eventual metastatic tumor. Therefore, we tested whether human PDAC metastases show evidences of enrichment for our gene signature compared to the primary tumor. To do so, we utilized singscore^86^ which uses rank-based statistics to score a sample’s gene expression profile with respect to a gene signature. The gene signature can include both an up-regulated set and a down-regulated set. We identified the human orthologs of our metastasis-high genes (n = 202) and metastasis-low genes (n = 174) and used them respectively as the up-set and the down-set for singscore. We call this version of singscore the MetScore. We observed that the MetScore is significantly higher in metastatic tumors compared to primary tumors in both PACA-CA and PACA-US cohorts (two-sided Wilcoxon rank-sum test, *p*-value 0.040 for PACA-CA and 9.1E-6 for PACA-US) (**Fig. 7A**). The large sample sizes in PACA-US provided adequate statistical power to perform a primary to metastasis comparison stratified by classical/basal classification. Across both molecular subtypes, metastases demonstrated higher average MetScores than primary tumors (*p*-value 4.1E-4 for classical and 0.048 for basal), suggesting that our gene signature is relevant to both classical and basal PDAC despite being borne out of an analysis of classic subclones only. Furthermore, the difference in MetScores between primary tumors and metastases was still apparent when we isolated the rapid autopsy patients in the PACA-US cohort harboring both primary tumors and metastases at the time of sampling and performed a paired intra-patient comparison (linear mixed effects model, *p-*value 4.5E-4; **Supp. Fig. 3A**), thereby eliminating any potential confounding relating to inter-patient differences. MetScores appeared to be similar across anatomic sites in the two cohorts (**Supp. Fig. 3C**), suggesting that our signature is associated with a generalized metastatic advantage rather than tropism to a specific anatomic site. These results suggest that our bi-directional gene signature identified in the mouse is also associated with metastatic potential in human PDAC.

**Figure 7.**
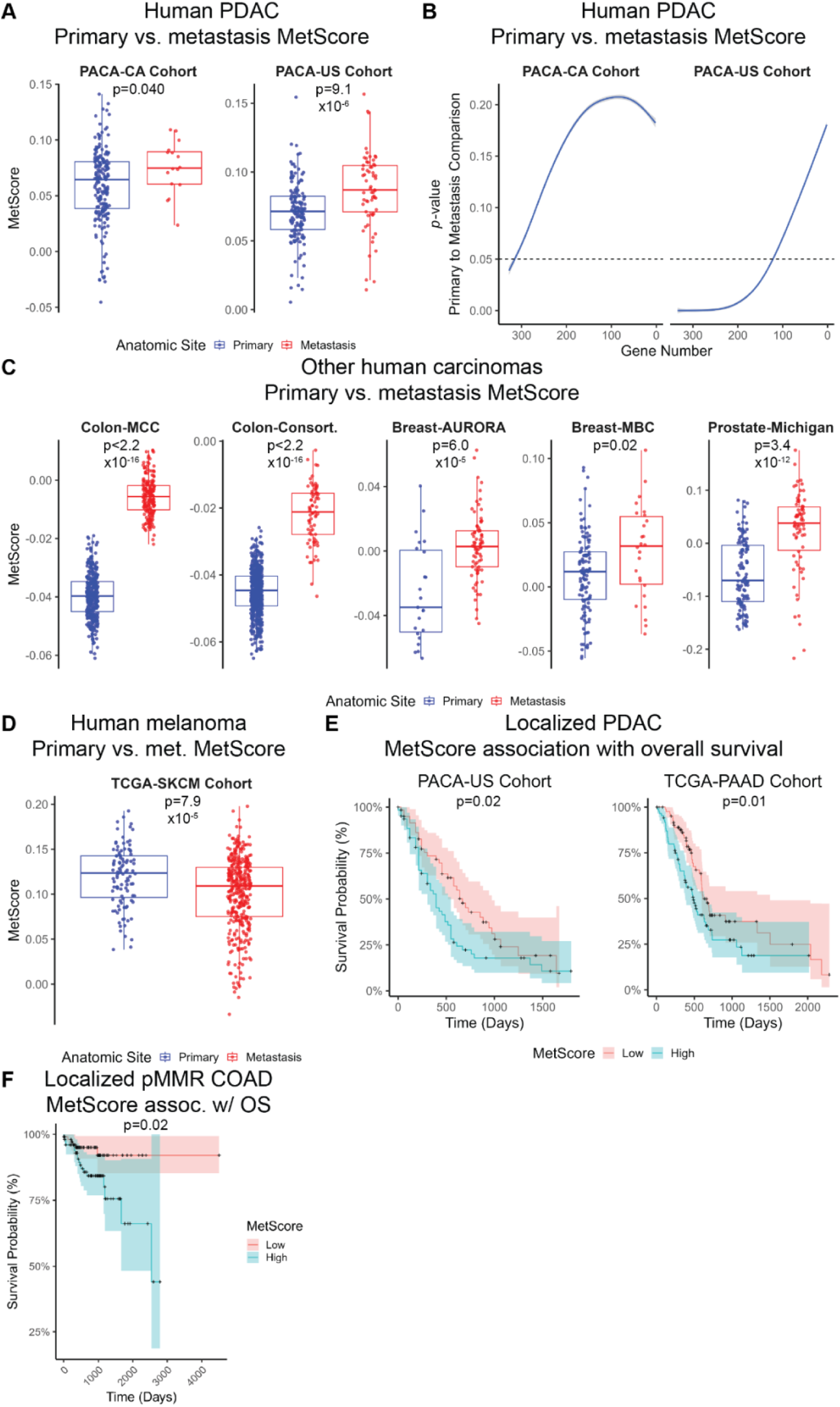
Metastasis-high and metastasis-low genes define a metastasis signature in human carcinomas. (**A, C-D**) Box plots depicting distributions of MetScores with overlaid points representing individual tumors for primary and metastatic samples in the indicated human patient datasets. *p*-values calculated using two-sided Wilcoxon rank sum tests. (**B**) Generalized additive model trendlines with 95% confidence intervals for *p*-values for primary to metastasis MetScore comparisons as a function of the number of genes from the complete metastasis-high and metastasis-low gene sets utilized. For each gene number, 1000 random samplings from the starting gene sets were performed. For each random draw, MetScores were calculated for all samples in the indicated cohort, primary and metastatic samples were compared, and a two-sided Wilcoxon rank sum test was used to generate a *p*-value. Random draws with zero genes in either the upregulated set or the downregulated set were discarded and not replaced. The *p*-values for all random draws at all tested gene numbers were used to generate the shown trendline. (**E-F**) Kaplan-Meier curves depicting overall survival for patients in the noted datasets stratified by MetScore. Patients in each dataset were ranked by MetScore, the top half of which being considered “High” and the bottom half considered “Low”. *p*-values were calculated using log-rank tests.

To evaluate whether the entire metastasis-high and metastasis-low gene sets are needed for MetScore to differentiate between primary and metastatic PDAC tumors or conversely whether its discriminative ability is largely dependent on the performance of a handful of highly informative genes, we assessed MetScore’s ability to discriminate between primary and metastatic PDAC with increasing numbers of genes randomly removed from the up and down sets. In both cohorts, discriminatory power, as measured with a bootstrap test that computes the average *p*-value for primary to metastasis MetScore comparisons across 1000 random samplings from the up and down sets for that total gene number, diminished immediately and rapidly as gene number decreased (**Fig. 7B**). When using PACA-CA as our test data, the smaller of the two cohorts, MetScore was particularly sensitive to decreases in gene number and required above 315 genes to consistently reach statistical significance. When using PACA-US as our test data, a larger cohort, MetScore was more robust, requiring above only 124 genes to consistently reach statistical significance. These results are consistent with metastatic potential being driven by the collective contributions of a large number of genes.

The steps of the metastatic cascade are similar for different cancer subtypes; we thus wondered whether the factors that determine metastatic potential are conserved across subtypes. We found that MetScores were significantly higher in metastases compared to primary tumors in all carcinoma cohorts we tested, including two colon adenocarcinoma (COAD) cohorts (two-sided Wilcoxon rank-sum test, *p*-value < 1E-16 for Colon-MCC^87^ and < 1E-16 for Colon-Consortium^87^), two breast invasive carcinoma (BRCA) cohorts (*p*-value 6.0E-5 for Breast-AURORA-US^88^ and 0.018 for Breast-MBC^89^), and one prostate adenocarcinoma (PRAD) cohort^90^ (*p*-value 3.4E-12; **Fig. 7C**). These results suggest that similar mechanisms underlie metastatic potential in a broad group of carcinomas, which are all epithelial in origin. In contrast to carcinomas, we observed an inversed trend in a melanoma cohort^91^with primary tumors having significantly higher MetScores than metastases (*p*-value 7.9E-5) (**Fig. 7D**). Melanoma is non-epithelial in origin^92^. Therefore, the gene signature associated with metastatic potential in mouse PDAC is enriched specifically in metastases of human carcinomas. These results suggest that similar factors underlie metastatic potential in carcinomas but are not broadly shared with melanoma.

### MetScore is associated with overall survival in localized human PDAC and pMMR COAD

We reasoned that localized PDAC patients whose primary tumors have a higher MetScore are likely to have worse overall survival because heterogenous primary tumors with a greater fraction of subclones in a metastasis-high state would likely engender distant recurrence earlier. We analyzed the survival curves of localized PDAC patients in two cohorts that have primary tumor transcriptomic data available. One cohort includes localized PDAC patients isolated from the previously-analyzed PACA-US cohort and the second is The Cancer Genome Atlas’s (TCGA’s) Pancreatic Adenocarcinoma (TCGA-PAAD) cohort^93^. In each cohort, we compared the overall survival of patients with tumors in the top half of MetScores to those in the bottom half (**Fig. 7E**). Multiple tumor properties other than metastatic potential contribute to overall survival, including therapy resistance, ability to induce cachexia, and thrombogenicity. Despite these complexities, the patients in the top half of tumor MetScores had a significantly worse overall survival in both cohorts (Log rank test, *p*-value 0.02 for PACA-US and 0.01 for TCGA-PAAD). Similarly, patients with localized mismatch repair proficient (pMMR) COAD in the TCGA-COAD cohort^94^ in the top half of tumor MetScores had significantly worse overall survival (*p*-value 0.02; **Fig. 7F**). We could not explore the relationship between MetScore and overall survival for the localized mismatch repair deficient (dMMR) COAD patients in the TCGA-COAD cohort because none of those patients died during the study period. There was not a statistically significant association between primary tumor MetScore and overall survival for patients with ductal or lobular histologies of BRCA in the TCGA-BRCA cohort^95^ or patients with localized PRAD in the TCGA-PRAD^96^ cohort or melanoma in the TCGA-SKCM cohort (**Supp. Fig. 3C**), which is expected given that initial metastatic potential is less likely than drug resistance to be the major determinant of patient outcome for these cancer subtypes in which certain patients have robust and long-lasting therapeutic responses. Overall, these results demonstrate the clinical relevance of our findings which provide a pathway for estimating how rapidly a patient with an early-stage pre-metastatic carcinoma is expected to develop distant metastases.

## Discussion

In the present study, we used DNA barcoding to uniquely label heterogeneous subclones in mouse primary PDAC tumors and measure their performance in metastasis assays. We observed a striking bimodality in the metastatic potential of subclones, with only a subset having high metastatic potential. We then analyzed open chromatin and transcriptome in each subclone in its pre-metastatic state to understand the molecular features associated with its metastatic potential. This analysis identified a set of genes enriched in subclones with a high metastatic potential and another set depleted in these subclones, a combination we call the metastatic gene signature. This gene signature identified IL-1 pathway genes with NF-κB and mesenchymal transcription factor activity resulting in a higher metastatic potential while genes involved in neuroendocrine differentiation, motility, and the canonical and non-canonical Wnt pathways with CDX2 and HOXA13 activity were associated with lower metastatic potential. This bi-directional gene signature is enriched in a broad set of human carcinoma metastases and has prognostic value in pancreatic ductal adenocarcinoma and mismatch repair proficient colon adenocarcinoma, predicting patient survival.

This work establishes an approach for linking the molecular state of a cancer cell to its future fate and behavior in the metastatic cascade by combining DNA barcoding and *in vivo* competition assays. The unique complexities of metastasis make standard fate mapping approaches challenging to implement. It is not possible to track the thousands of cells that leave the primary tumor each day using live imaging. Even if imaging was possible, high-throughput molecular characterization of these single cells would be a formidable challenge. Moreover, because each tumor develops in a different and unique fashion, time course sampling from different tumors will not be informative. Our approach, in essence, captures cells in a mid-metastasis state, barcodes them, expands them for molecular characterization, then returns them to the mid-metastatic position to compare their performance. The competitive nature of these assays is a key accurate quantification of metastatic potential, allowing us to compare different cells in the exact same condition. Indeed, we observed high variation in overall metastatic tumor burden between different mice (data not shown) which would have easily masked clone-to-clone differences had we tested each clone separately. This approach builds on a strong body of literature in tracking metastatic performance of cancer cells using DNA barcodes^12–16,97–99^. Prior applications of DNA barcoding for studying clonal dynamics during PDAC metastasis necessitated the use of immunocompromised hosts^12,16^. In contrast, our models permit interrogation of metastatic mechanisms that rely on interactions with the host immune system. Furthermore, our approach is readily translatable to other cancers as well as processes beyond metastasis, such as drug resistance.

Placing metastasis-high and metastasis-low subclones in the natural history of PDAC development was insightful. The metastasis-high and metastasis-low subclones occupied a similar position along the normal-to-PDAC developmental trajectory and all belonged to the classical subtype. These observations run counter to a prevailing theory of PDAC development which predicts highly metastatic subclones as being the most advanced along a normal-to-PDAC axis^100^. We speculate that the metastasis-high and -low states represent a branch at the terminus of the normal-to-PDAC axis, both with a high fitness in the primary tumor, but one far more adept at navigating the metastatic cascade. Accordingly, the majority of the metastasis-high and metastasis-low specific genes are upregulated in PDAC cells compared to normal; they are activated to a greater or lesser degree in one state versus the other.

Among the genes that are associated with a higher metastatic potential, an inflammation program stands out. A well-established link exists between inflammation and PDAC tumorigenesis, with pancreatitis being a significant clinical risk factor for PDAC^101^ and necessary for tumorigenesis in *Kras*-driven mouse models^102^. Furthermore, previous studies have suggested that inflammation promotes PDAC metastasis^103,104^. However, progress towards targeting inflammation in PDAC patients has stalled. Clinical trials of broad anti-inflammatory drugs in advanced PDAC patients have been disappointing^105,106^, likely because inflammatory pathways are active not only in cancer cells but in the many stromal cells present in the tumor microenvironment leading to pleiotropic effects. We are encouraged by the discovery of two downstream transcriptional targets of NF-κB, *Fos* and *Il23a*, as being novel functional mediators of PDAC metastasis whose inhibition may elicit a more cancer cell-specific response. While c-Fos, the AP-1 family transcription factor encoded by *Fos*, has been implicated in PDAC pathogenesis^107,108^, our experimental design allowed us to detect a specific contribution for c-Fos to liver metastasis. The pro-inflammatory cytokine IL-23α, encoded by *Il23a*, has been demonstrated to promote metastasis of other cancer subtypes^109,110^ but its role in PDAC metastasis has been unclear. Both c-Fos and IL-23α have specific inhibitors^111–113^ in clinical use that can be advanced to pre-clinical testing. The 196 genes unrelated to inflammation upregulated in subclones with a high metastatic potential did not associate as clearly with a single pathway, pointing to the complexity of metastatic potential. We validated two of these additional met-high genes, *Myo1b* and *Tmem40*, as being novel functional regulators of PDAC metastasis. While *Myo1b* has been shown to promote metastasis of other cancer subtypes^114,115^, we believe we are the first to demonstrate that *Tmem40* is a functional mediator. Their respective gene products, MYO1B and TMEM40, can be advanced to drug development while larger screens should be performed to evaluate the functional impact of all the remaining metastasis-high genes on metastatic potential.

Our dataset highlighted the activation of developmental transcription factors as a prominent feature of the metastasis-low state. These included transcription factors related to islet specification and development (e.g., ISL1, PBX1, MNX1, and FOXP1) and the ParaHox transcription factor CDX2, which differentiates developing post-gastric gut endoderm, giving rise to the pancreas, from pre-gastric. These findings are consistent with the developmental constraint model of cancer evolution in which cancer cells evolve by accessing gene programs specific to developmental lineages adjacent to the cell of origin^56^. However, the Hox family TF HOXA13, which is strongly activated in the metastasis-low subclones, is expressed in developing mesoderm and ectoderm for anterior-posterior patterning of the musculoskeletal and nervous systems and not endoderm, which gives rise to pancreatic epithelium^78^. This observation points to a complex multi-lineage state involving extreme dedifferentiation, which would broaden the constraints in the developmental constraint model. The accentuated activation of developmental transcription factors in metastasis-low subclones suggests their positive selection in the primary tumor, while their deemphasis in metastasis-high subclones implies their dispensability or detriment for metastasis. Future studies are needed to delineate the specific effects of each developmental program in each tumor compartment.

To explore the relevance of our findings in mouse models to human PDAC biology, we developed the MetScore, which produces single sample numerical scores based on enrichment and depletion of our identified metastasis-high and metastasis-low genes, respectively, in bulk transcriptomic data. We observed that human PDAC metastases have higher MetScores than primary tumors and that localized PDAC patients whose primary tumors have higher MetScores have worse overall survival, indicating that the signature we discovered in mice is at least partially conserved in humans. Remarkably, in all other human carcinomas evaluated, metastases similarly had higher MetScores than primary tumors, suggesting this signature may also be conserved across epithelial cancer subtypes. These results suggest that the MetScore may be a useful tool for predicting metastasis risk for patients with localized carcinomas. This would provide critical prognostic information to patients, empowering them to make better informed decisions about their care, and may also serve as a predictive biomarker for the treating oncologist in cases where the optimal intensity of adjuvant therapy is unclear (e.g., stage II colon adenocarcinoma). The metastasis risk prediction tools OncotypeDX®^116^ and MammaPrint®^117^ have revolutionized the care of early-stage breast cancer by allowing oncologists to de-escalate the intensity of adjuvant therapy for women with a low risk of distant recurrence, thereby preventing overtreatment. We envision MetScore having a similar impact for other carcinoma subtypes where decisions regarding the intensity of adjuvant therapy are currently made using clinical risk factors alone. Furthermore, in contrast to the OncotypeDX® and MammaPrint®, MetScore utilizes a large number of genes (376 total as opposed to 21 for OncotypeDX® and 70 for MammaPrint®) and generates its score based on not only enrichment of met-high genes but also depletion of met-low genes, which we predict will make MetScore an especially robust scoring metric.

The limitations of this study include the number of subclones and tumors which may be small relative to the diversity of metastatic states that could exist within a tumor and across tumors. However, the correlations and prognostic prediction in humans indicate that the signature identified here captures an important fraction of genes associated with metastatic potential. Capturing the full diversity of highly metastatic states in the future is likely to provide a more comprehensive set of metastasis-associated genes and refine the present set.

Overall, this study describes a new approach for linking cell state to future cellular behavior and provides important insights into the underlying biology of PDAC metastasis, laying the groundwork for novel therapeutic strategies and diagnostic tools and offering hope for improved outcomes for patients with this devastating disease.

## Methods

### Cell Lines

The KPC-1 and KPC-2 cell lines were a gift from Dr. Lei Zheng. They were generated from primary tumors of KPC mice^21^ (i.e., *Pdx1-Cre;LSL-Kras^G12D/+^;Trp53^R^*^172^*^H/+^*) as described previously^22,23^. Cell lines were tested for mycoplasma contamination using a PCR-based kit (Bulldog Bio 2523348) and were found to be mycoplasma negative.

### Barcoding

The DNA barcodes utilized in this study were based on a previously published homing guide RNA (hgRNA) library^24,25^. Two transposable plasmid libraries with random bases were mixed. The first library contains two stretches of degenerate bases, one 15 bases in length (“LeftBarcode”) and the other 10 bases in length (“RightBarcode”). These two stretches are separated by a constant 94 base region. It was constructed from Addgene #104536 plasmid as described previously^24^. The second library is very similar. It contains two stretches of degenerate bases, one 19 bases in length (“LeftBarcode”) and the other 10 bases in length (“RightBarcode”) that are separated by a constant 94 base region; however, it also contains a puromycin resistance marker expressed by the EF-1α promoter. It was constructed from Addgene #104537 plasmid as described previously^24^. The inserts in both plasmid libraries are flanked by PiggyBac inverted repeats which enables their integration into the genome using the PiggyBac transposase. The constructs express the barcodes in small RNA form from a U6 promoter. The universal amplification primers for these barcodes allow reliable identification in sequencing based upon either the forward or reverse reads.

Nucleofection was used to introduce the barcode libraries into KPC cells (Lonza Basic Nucleofector^TM^ Kit for Primary Mammalian Epithelial Cells [VPI-1005] and Nucleofector® II Device using program T-020). Barcode libraries were co-transfected with Super PiggyBac Transposase Expression Vector (System Biosciences PB210PA-1) to facilitate integration of the barcode construct into the genome. Two strategies were utilized to encourage a large number of barcode insertions per cell. The first strategy was co-transfection of the *Ins21* and *Ins25-puro* libraries in a 19:1 ratio. Under these conditions, only cells with a large number of integrations would be likely to have integrated a puromycin resistance gene containing construct, allowing us to eliminate cells with few integrations during antibiotic selection. The second strategy was using a transposase:transposon ratio of 1:10 rather than the more typically used 1:3. Because PiggyBac can both integrate and excise transposons, having less PiggyBac in the cells reduces the likelihood that barcode constructs integrated into the genome will be removed during the initial transposition process.

Following barcode integration, the cells underwent puromycin (InvivoGen ant-pr-1) selection for seven days. Puromycin resistant cells were then sorted as single cells into wells of a 96 well plate using a Sony Sorter SH800. Propidium iodide (Invitrogen 00-6990-50) was used to exclude dead cells. The resulting colonies were expanded over the course of several weeks while remaining under puromycin selection and then cryopreserved in fetal bovine serum (Thermo Fisher 16140071) with 10% DMSO (Sigma-Aldrich D2650).

### Barcode sequencing – library preparation and sequencing

Genomic DNA was isolated from cells or mouse tumors using a DNeasy Blood and Tissue kit (Qiagen 69504) as per the manufacturer’s instructions. The hgRNA locus was amplified and sequenced using next generation sequencing as described previously^25^. Briefly, the hgRNA locus was amplified using primers with overhangs containing primer binding sites for Illumina sequencing by synthesis (i.e., PCR1). Then, a second PCR amplification was performed using primers with overhangs containing either P5 or P7 to facilitate binding to the Illumina flow cell and a random DNA sequence (i.e., the i5 or i7 index sequence) to facilitate pooling and deconvolution of multiple samples in the same sequencing run (i.e., PCR2). Deviating from the published protocol, a set of four forward and four reverse degenerate PCR1 primers were used to increase library diversity and custom PCR2 indexing primers were used to allow for pooling of a large number of samples (**Supp. Table 1**). The resulting libraries were then pooled, purified using a DNA Clean and Concentrator-5 kit (Zymo D4004), and quantified using either a Qubit® dsDNA HS Assay Kits (Thermo Fisher Q32851) or an NEBNext® Library Quant Kit for Illumina® (NEB E7630). The final libraries were sequenced using an Illumina MiSeq device and MiSeq Reagent Kit v2 (Illumina MS-102-2002).

### Barcode sequencing – data processing and analysis

Raw sequencing data was processed on a high-performance computing cluster (The Advanced Research Computing at Hopkins [ARCH] “Rockfish” cluster [https://www.arch.jhu.edu/about-arch/]) using the previously published pipeline^25^. Briefly, this pipeline decompresses the raw sequencing data, compiles Read 1 and Read 2 sequences from each sample to a list of paired LeftBarcodes and RightBarcodes, sequentially corrects for sequencing errors in the LeftBarcode and RightBarcode regions, and then compiles complete lists of LeftBarcode-RightBarcode counts for each sample.

To characterize the barcodes specific to each subclone, local Python scripts were used to first filter out unique LeftBarcode-RightBarcode pairs with fewer than 3-4 reads depending on sequencing depth. Then the RightBarcode sequences specific to that subclone were defined by filtering out RightBarcodes whose reads made up less than one percent of the total reads. Finally, each RightBarcode’s LeftBarcode mate was defined as the most abundant LeftBarcode out of those paired with that RightBarcode.

To characterize the subclones present in a tumor or *in vitro* culture, local Python scripts were used to first filter out unique LeftBarcode-RightBarcode pairs with fewer than three reads. Then, the identifier-spacer pairs found in that tumor were cross-referenced against LeftBarcode-RightBarcode pairs specific to each subclone included in the experiment (i.e., either all KPC-1 derived subclones or all KPC-2 derived subclones). Since all LeftBarcode-RightBarcode pairs were unique to their assigned subclone, the presence of a single LeftBarcode-RightBarcode pair was sufficient to indicate the present of its assigned subclone in the tumor. Using this strategy, each tumor was noted for the presence or absence of each subclone included in the experiment.

### Mouse models

All animal procedures were approved by Johns Hopkins University’s Animal Care and Use Committee (ACUC). C57BL/6J mice were obtained from The Jackson Laboratory (strain #000664). Eight-week-old female mice were used. Splenic, intraperitoneal, and orthotopic injections were performed as described previously^22,23,26,118^. For splenic and intraperitoneal injections, 500,000 cells were injected. For orthotopic injections, 50,000 cells were injected. In all cases, the injected animals were allowed to incubate for 4 weeks prior to sacrifice. Tumors were harvested in all cases with microdissection using a dissection microscope (ZEISS SteREO Discovery.V8).

### Gross pathology and histology

Photographs of mouse tumors were taken using an iPhone 13.

Mouse tumors were fixed in 10% neutral buffered formalin for 48 hours, after which they were processed into paraffin embedded tissue blocks as described previously^118^. They were then sectioned and stained with hematoxylin and eosin (H&E) as described previously^118^. Light micrographs of the H&E-stained sections were captured using a Zeiss Axio Vert.A1 microscope.

### Proliferation assays

KPC-1 and KPC-2 derived monoclonal lines were seeded onto wells of a 96 well plate (5,000 cells for KPC-1 and 10,000 cells for KPC-2). At 24 and 48 hours, culture medium was aspirated and the cells were placed at −70°C. Relative cell number at each timepoint was quantified using a CyQUANT Cell Proliferation Assay, for cells in culture (Thermo Fisher C7026), as per the manufacturer’s instructions.

### Genotyping

Genomic DNA was isolated from each KPC-1 and KPC-2 derived monoclonal line using a DNeasy Blood and Tissue kit (Qiagen 69504) as per the manufacturer’s instructions. Genotyping PCR reactions were performed according to The Jackson Laboratory for *Pdx1-Cre*^119^, and the Tyler Jacks Laboratory for *Lsl-Kras^G12D^* ^120^ and for *Lsl-Trp53^R17H^* ^121^. PCR products were separated on a 1% agarose gel run at 8V/cm for one hour and visualized with SYBR Gold reagent (Thermo Fisher S11494) as per the manufacturer’s recommendations.

### ATAC-seq – library preparation

ATAC-seq libraries were generated from 50,000 cryopreserved cells harvested from *in vitro* cultures as described previously^122^. Libraries were then pooled and sequenced on a Novoseq 6000 using 100 bp paired end reads.

### ATAC-seq – data processing and analysis

#### Data processing

Raw sequencing data was processed on a high-performance computing cluster (the Rockfish cluster described in Barcode sequencing – data processing and analysis) using the ENCODE project’s publicly available ATAC-seq pipeline^123^ with default settings. Briefly, adapters were trimmed using cutadapt 1.9.1, trimmed reads were aligned to the mm10 genome using Bowtie2^124^ 2.2.6, low quality, mitochondrial, and duplicate reads were filtered out using SAMtools^125^ 1.7 with Picard^126^ 1.126 to mark duplicates, and peaks were called using Macs2^127^ 2.1.0. For two samples, KPC-2_LoC and KPC-2_HiA, two independent libraries were prepared from separate aliquots of cryopreserved cells (i.e., technical replicates). Raw sequencing data from technical replicates was pooled prior to processing.

ATAC-seq data from normal, pre-neoplasia, pancreatitis, pre-neoplasia+pancreatitis, and primary PDAC generated by Alonso-Curbelo and colleagues^28^ was downloaded from NCBI (BioProject #PRJNA548087) and processed in the same manner.

Locally in R, a consensus peak set was generated by merging overlapping peaks from the individual samples’ peak sets using DiffBind^128^ 3.14.1. Then, a normalized count matrix, i.e., the number of reads aligning to each peak, normalized by the total reads in peaks for that sample, across all samples, was generated using DiffBind.

#### Batch correction

For analyses that included both the metastasis-high and metastasis-low subclones generated in this study as well as the normal, pre-neoplasia, pancreatitis, pre-neoplasia+pancreatitis, and primary PDAC generated by Alonso-Curbelo and colleagues^28^, the normalized count matrix was batch corrected using ComBat^29^ from the sva package (version 3.52.0). A parametric batch correction was performed with sample type (i.e., normal, pancreatitis, pre-neoplasia, pre-neoplasia+pancreatitis, PDAC) included as a covariate and with the Alonso-Curbelo *et al*. samples defined as the reference batch.

#### Principal component analysis

Principal component analysis was performed using the lo*g*_2_(*x* + 1) transformed and scaled normalized count matrix using base R’s prcomp() function.

#### Differential accessibility analysis

The generalized linear model functionality of DESeq2^35^ was utilized to identify differentially accessible peaks between metastasis-high and metastasis-low subclones while controlling for parental group status by modeling the metastatic potential (i.e., high vs. low) as a fixed effect and parental group (i.e., KPC-1 vs. KPC-2) as a random effect (`design: ∼parental_group + metastatic_potential`). An FDR cutoff of 0.05 was used.

#### Peak profile plot

To visualize ATAC-seq signal profiles for peaks with increased accessibility in metastasis-high subclones and separately for peaks with increased accessibility in metastasis-low subclones across all of the tested subclones, a matrix containing scores for the genomic regions of interest was first generated using the plotProfile() function of DiffBind 3.14.1 and then plots were generated using the plotProfile command of deeptools^129^ 3.5.5.

#### Peak annotation

Significantly differentially accessible peaks were assigned to their nearest genes using the `annotatePeak()` function from ChIPseeker^130^ 1.40.0 with the mm10 genome as the reference genome. In addition to identifying the nearest gene, this function also assigned the peaks to one of several bins based on its distance to the nearest gene’s transcription start site (TSS). For visualization purposes, the “Distal Intergenic” and “Downstream (<=300bp)” categories were merged into a “Distal Intergenic” category and the “Promoter (<=1kb)”, “Promoter (1-2kb)”, and “Promoter (2-3kb”) categories were merged into a “Promoter” category (**Fig. 3E**). All mouse candidate *cis*-regulatory elements (CREs; mm10 genome) identified by the ENCODE project were downloaded from SCREEN: Search Candidate cis-Regulatory Elements by ENCODE Registry of cCREs V3^131^. First, peaks were assessed for overlap with any identified candidate CRE. Then, overlaps were broken down by category. For visualization purposes, “pELS” and “dELS” categories were merged into “Enhancer”; “PLS” and “CA-H3K4me3” categories were merged into “Promoter”; and “CA”, “CA-TF” and “TF” were merged into “Candidate RE, NOS” (**Fig. 3F**).

#### Analysis of peak enrichment within regulatory regions corresponding to genes in GO^37,38^ (Gene Ontology) Biological Process gene sets

Metastasis-high and metastasis-low peaks were separately inputted into the GREAT^39^ (Genomic Regions Enrichment of Annotation Tools) web portal using the mm10 genome as the reference genome. Default settings were used. All enriched pathways reported here passed an FDR threshold of 0.05 for both binomial and hypergeometric tests.

### RNA-seq – library preparation

Total RNA was isolated from KPC cells being grown *in vitro* by performing a TRizol (Thermo Fisher 15596026) chloroform (Sigma-Aldrich C2432) extraction followed by cleanup using an RNA Clean and Concentrator-5 kit (Zymo R1013) as per the manufacturer’s instructions. mRNA was then isolated using a NEBNext Poly(A) mRNA Magnetic Isolation Module (New England Biolabs E7490). Libraries were prepared using an xGen RNA Library Prep Kit (IDT 10009814) and xGen UDI indexing primers (IDT 10005922) according to the manufacturer’s instructions. The final libraries were sequenced on a Novoseq 6000 using 150 bp paired end reads.

### RNA-seq – data processing and analysis

#### Data processing

Raw RNA-seq data was processed on a high-performance computing cluster (the Rockfish cluster described in Barcode sequencing – data processing and analysis). Adapter sequences were trimmed using Trimmomatic^132^ 0.39. Transcripts were then quantified using Salmon^133^ 1.10.1 in the mapping-based mode with GC bias correction. The M33 (GRCm39) transcript sequences from GENCODE^134^ were used as the reference transcriptome.

RNA-seq data from normal, pre-neoplasia, pancreatitis, pre-neoplasia+pancreatitis, and primary PDAC generated by Alonso-Curbelo and colleagues^28^ was downloaded from NCBI (BioProject PRJNA547612) and processed in the same manner.

Then, locally in R, transcript quantifications were imported and summarized to the gene level using Tximeta^135^ 1.22.1.

#### Differential expression analysis

For the metastasis-high vs. metastasis-low comparison, genes with fewer than 50 reads across all KPC samples were first excluded. Then, the generalized linear model functionality of DESeq2 1.44.0 was utilized to identify differentially expressed genes between metastasis-high and metastasis-low subclones while controlling for parental group status by modeling the metastatic potential (i.e., high vs. low) as a fixed effect and parental group (i.e., KPC-1 vs. KPC-2) as a random effect (`design: ∼parental_group + metastatic_potential`). An FDR cutoff of 0.05 was used.

For the primary PDAC vs. normal pancreas comparison, genes with fewer than 100 reads across all KPC and *in vivo* samples were first excluded. Then, DESeq2 was used to identify differentially expressed genes between primary PDAC samples and normal pancreas samples. An FDR cutoff of 0.05 was used.

#### Classical/basal subtype classification

To classify the KPC subclones as being either classical or basal subtype, the PurIST classifier^34^ was used to calculate the probability of basal subtype classification for each sample. This was performed in R by following the publisher’s instructions. The human classical and basal subtype defining genes used by the classifier were converted to their mouse orthologs using the Ensembl^136^ BioMart web portal. Bulk RNA-seq data from each subclone was used as input to the classifier, specifically gene TPM (transcripts per million; see RNA-seq – library preparation and RNA-seq – data processing and analysis).

### Signal tracks

Since the raw RNA-seq data was initially processed using a pseudoalignment method (Salmon), it was re-processed here as aligned reads are needed for visualization as a signal track.

Adapter trimmed RNA-seq reads were aligned to the mm10 genome using STAR^137^ 2.7.10a with default settings.

Aligned reads, ATAC-seq or RNA-seq, for each sample were converted to bigWig track format using `bamCoverage` from deeptools 3.5.5. A bin size of 25 bp was used and signals were normalized by total library size (RPKM; reads per kilobase per million mapped reads). Average bigWig tracks were then generated for the metastasis-high and metastasis-low groups of samples using the `mean` command from the WiggleTools^138^ package (version 1.2) followed by conversion of the resulting Wig formatted tracks back to bigWig format using the `wigToBigWig` command from the UCSC Genome Browser^139^ suite of command-line utilities.

Signal tracks were then plotted using CoolBox^140^ 0.3.8.

All of the above steps were performed on a high-performance computing cluster (the Rockfish cluster described in Barcode sequencing – data processing and analysis).

### Analysis of gene set enrichment amongst metastasis-high and metastasis-low genes

Enrichment of GO^37,38^ and KEGG^40^ (Kyoto Encyclopedia of Genes and Genomes) pathways amongst metastasis-high and metastasis-low genes was assessed in R using the `enrichGO()` and `enrichKEGG()` functions in clusterProfiler^141^ 4.12.0. An FDR cutoff of 0.05 was used.

### Transcription factor footprinting

Non-redundant vertebrate transcription factor binding profiles, i.e., motifs, were downloaded from the JASPAR database^77^. Locally in R, this set of motifs was then filtered to exclude motifs corresponding to non-expressed or lowly expressed transcription factors in the KPC cells. Non-expressed genes were defined as those having fewer than 50 total reads across all samples in our RNA-seq dataset and lowly expressed genes were defined as those whose statistical significance was not calculated by DESeq2 due to low expression in our differential expression analysis.

Then, on a high-performance computing cluster (the Rockfish cluster described in Barcode sequencing – data processing and analysis), aligned ATAC-seq reads for every KPC sample were downsampled to the coverage of the least covered sample using the `view` command in samtools 1.15.1. Then, aligned ATAC-seq reads were merged amongst the metastasis-high samples and separately amongst the metastasis-low samples using the `merge` command in samtools 1.15.1.

Finally, on the Rockfish cluster, transcription factor footprinting was performed by inputting the filtered motifs (n = 511), the metastasis-high and metastasis-low merged aligned ATAC-seq reads, and the complete consensus peak set for all KPC samples (n = 176,964 peaks) to TOBIAS^76^ 0.15.1. The `ATACorrect` command was used to correct the ATAC-seq signal in the merged samples for Tn5 insertion bias. Then, the `ScoreBigwig` command was used to calculate a continuous footprint score across peaks in the consensus peak set based on the depletion of signal and the general accessibility of the nearby region. Finally, the `BINDetect` command was used to (1) identify putative TF binding sites within peak regions by matching the known motifs to the peak region DNA sequences using MOODS^142^ (MOtif Occurrence Detection Suite); (2) classify each putative TF binding site as being either bound or unbound in each condition (i.e., metastasis-high and metastasis-low) based on a footprint score cutoff; (3) calculate the *log*_2_ fold change in footprint score between the two conditions for each binding site; (4) calculate a differential binding score (DBS) for each motif representing the global distribution of *log*_2_ fold changes across binding sites for that motif; and (5) calculate a *p*-value for each motif by comparing its DBS to 100 DBSs generated from randomly sampled *log*_2_ fold changes from the background distribution.

Locally in R, differential binding score *p*-values were adjusted for multiple hypothesis testing using a Bonferroni adjustment. An adjusted *p*-value cutoff of 0.05 was used.

### Targeted shRNA screen

#### shRNA design

We designed three miR-30-based shRNAs targeting each gene included in the screen as well as three negative control shRNAs targeting luciferase, which the KPC cells do not express. The majority of the shRNA sequences used were designed using the SplashRNA algorithm^143^. For genes for which the SplashRNA algorithm did not produce at least three shRNA sequences with SplashRNA scores greater than one, the remaining sequences were selected from the shRNAs in Table 3 in Fellmann et al., 2013^144^. All shRNA sequences used in this study can be found in **Supplementary Table 2**.

#### Cloning and lentiviral transduction

shRNAs were cloned in a pooled fashion into the SGEP lentiviral expression vector as described previously^145^. SGEP was obtained from Addgene (#111170). The resulting shRNA library was packaged into lentiviral particles via co-transfection with VSV.G (Addgene #12259) and gag/pol (Addgene #12263) plasmids into HEK293T cells (ATCC CRL-3216) using Lipofectamine^TM^ 2000 Transfection Reagent (Thermo Fisher 11668027). Lentiviral particles were concentrated via precipitation with lentiviral concentration solution (4X stock is 40% [W/V] PEG-8000 and 1.2M NaCl in PBS [pH 7]). KPC cells were transduced in the presence of polybrene using a low multiplicity of infection. GFP positive cells were isolated using a Sony Sorter SH800S. Dead cells were excluded using propidium iodide (Thermo Fisher 00-6990-50).

#### shRNA amplicon sequencing

Genomic DNA was isolated from a day zero pre-injection sample and mouse tumors using a DNeasy Blood and Tissue Kit (Qiagen 69504) according to the manufacturer’s instructions. When tumors were too large to be digested and loaded onto a single spin column, they were subdivided into smaller chunks, each of which was digested and library prepped separately. Three independent draws of the day zero pre-injection sample were library prepped separately (i.e., three technical replicates). The shRNA locus was amplified using primers flanking the entire shRNA sequence with overhangs containing primer binding sites for Illumina sequencing by synthesis (i.e., PCR1; **Supp. Table 1**). Then, a second PCR amplification was performed using primers with overhangs containing either P5 or P7 to facilitate binding to the Illumina flow cell and a random DNA sequence (i.e., the i5 or i7 index sequence) to facilitate pooling and deconvolution of multiple samples in the same sequencing run (i.e., PCR2; **Supp. Table 1**). The resulting libraries were then pooled, purified using a DNA Clean and Concentrator-5 kit (Zymo D4004), and quantified using a NEBNext® Library Quant Kit for Illumina® (NEB E7630S). The final libraries were sequenced using an Ilumina MiSeq device and MiSeq Reagent Kit v2 (Illumina MS-102-2002).

#### Data processing and analysis

First, reads from each sample were mapped to the expected shRNA sequences present in the library. To accomplish this, Bowtie^146^ 1.3.0 was used to align the region of Read1 expected to correspond to the variable region of the shRNA-guide stem against a reference composed of the variable regions present in the library. One base pair mismatches were allowed to account for sequencing error. This produced a list of counts for each shRNA in the screen for each sample. This analysis was performed on a high-performance computing cluster (the Rockfish cluster described in Barcode sequencing – data processing and analysis).

Next, locally in R, shRNA counts for the three day zero pre-injection samples were combined to create a pooled day zero pre-injection sample. shRNAs whose abundance in the pooled day zero pre-injection sample was less than 0.25% were excluded from further analysis steps. Tumors with less than 100 shRNA counts were also excluded from further analysis steps.

The tumor samples were then normalized by dividing counts for each shRNA by the total shRNA counts for that respective sample. Normalized shRNA counts in samples derived from the same tumor (i.e., cases in which the tumor was too large to be digested and loaded onto a single spin column) were combined and the resulting pooled samples were re-normalized by dividing the normalized counts for each shRNA by the total normalized counts for that pooled sample.

Enrichment/depletion of shRNAs between liver metastases and primary tumors was evaluated as described previously^147^. Briefly, normalized shRNA counts were subjected to linear regression modeling with the sample type (i.e., primary tumor vs. liver metastasis) as a covariate using the `lm()` function in R. The resulting coefficients, standard errors (*SE*), *t*-values, and *p*-values were extracted for each shRNA. To account for variability and normalize the data, *log*_2_ fold changes of the non-targeting control shRNAs were normalized using the bestNormalize package (version 1.9.1) to create a transformation object. This transformation was then applied to the *log*_2_ fold changes of targeting shRNAs to calculate *z*-scores.

For weighted combined *p*-value calculation, *z*-scores were divided by the *SE*, and weights were defined as the inverse of the *SE* squared. Subsequently, shRNAs targeting the same gene were aggregated, and the adjusted *z*-scores were summed and normalized by the square root of the sum of the weights to generate combined *z*-scores for each gene. The combined *z*-scores were then used to calculate *p*-values by applying a two-tailed normal distribution test (2*pnorm(-abs(combined.z))). A *p*-value cutoff of 0.05 was used.

### Scoring human tumor samples for a metastatic gene signature (i.e., the MetScore)

The mouse metastasis-high (n = 207) and metastasis-low (n = 182) genes were converted to their human orthologs using the Ensembl^136^ BioMart web portal. Since not all of these genes have well characterized human orthologs, this produced slightly smaller sets of human metastasis-high (n = 202) and metastasis-low (n = 174) genes.

Bulk transcriptomic data corresponding to human primary tumors and metastases in the form of normalized signal (microarray) or normalized count (RNA-seq) matrices were obtained from the following sources:

| **<u>Cohort</u>** | **<u>Source</u>** |
| --- | --- |
| PACA-CA<sup>84,85</sup> | pdacR<sup>148</sup> |
| PACA-US<sup>31</sup> | pdacR<sup>148</sup> |
| TCGA-PAAD<sup>93</sup> | UCSC Xena Browser<sup>149</sup> |
| Colon-MCC<sup>87</sup> | Gene Expression Omnibus<sup>150</sup>, GSE131418 |
| Colon-Consortium<sup>87</sup> | Gene Expression Omnibus<sup>150</sup>, GSE131418 |
| Breast-AURORA-US<sup>88</sup> | Gene Expression Omnibus<sup>150</sup>, GSE193103 |
| Breast-MBC<sup>151</sup> | cBioPortal<sup>152-154</sup> |
| Prostate-Michigan<sup>90</sup> | Gene Expression Omnibus<sup>150</sup>, GSE35988 |
| TCGA-SKCM<sup>91</sup> | UCSC Xena Browser<sup>149</sup> |
| TCGA-COAD<sup>94</sup> | UCSC Xena Browser<sup>149</sup> |
| TCGA-BRCA<sup>95</sup> | UCSC Xena Browser<sup>149</sup> |
| TCGA-PRAD<sup>96</sup> | UCSC Xena Browser<sup>149</sup> |

For microarray datasets in which there were multiple probes per gene (Colon-MCC, Colon-Consortium, Prostate-Michigan), the probe with the greatest average signal for each gene was kept and the remaining probes for that gene were discarded.

For microarray datasets in which there were multiple probe sets utilized (Prostate-Michigan), only genes present in all probe sets were considered.

For RNA-seq datasets (Breast-MBC) in which there were multiple entries assigned to the same gene, presumably corresponding to different mRNA isoforms, the normalized counts for these independent entries were summed to create a single entry for each gene.

In certain cases, filtering to isolate tumors with the desired histologic subtype or molecular phenotype was required. From the TCGA-PAAD dataset, tumors with neuroendocrine histology were excluded. From the TCGA-COAD dataset, primary tumors with mismatch repair proficiency or deficiency were separately isolated and tumors lacking any annotation were excluded. From the TCGA-BRCA dataset, tumors with ductal (“Infiltrating duct carcinoma, NOS”) or lobular (‘‘“Lobular carcinoma, NOS”) histology were separately isolated.

For RNA-seq datasets, non- and lowly expressed genes were excluded (average FPKM < 1 for TCGA datasets and average TPM < 1 for PACA-CA, Breast-AURORA-US, and Breast-MBC).

To calculate the MetScore for each sample, the genes within that sample were first ranked using `rankGenes()` from singscore^86^ 1.24.0 in R. Then, the MetScore was calculated using `singscore()` with the human metastasis-high genes as the upSet and the human metastasis-low genes as the downSet. Notably, not all of our metastasis-high and metastasis-low genes were detected in each dataset after filtering out non- or lowly expressed genes. Only the detected metastasis-high and metastasis-low genes after filtering steps were considered.

Average MetScores were then compared between primary tumor and metastasis samples within each patient cohort. Statistical significance was determined using a two-sided Wilcoxon rank sum test. A *p*-value cutoff of 0.05 was used.

For one cohort, PACA-US, a subset of the rapid autopsy patients for which both primary tumor and metastasis samples were available was isolated (**Supp. Fig. 3A**). For this sub-cohort, MetScores were compared between primary tumor and metastasis samples while controlling for inter-patient differences by modeling anatomic site as a fixed effect and patient as a random effect in a linear mixed effects model. A *p*-value cutoff of 0.05 was used.

The relationship between primary tumor MetScore and overall survival was then explored. Primary tumor samples from localized cancer patients for which survival information was available were isolated from the PACA-US, TCGA-PAAD, TCGA-COAD, TCGA-BRCA, TCGA-PRAD, and TCGA-SKCM cohorts. Samples were stratified by MetScore with patients in the top half of their respective cohort being designated “high” and patients in the bottom half being designated “low”. Kaplan-Meier curves were generated using the survival^155,156^ 3.5-8 and ggfortify^157,158^ 0.4.17 packages in R. Statistical significance was determined using log rank tests with a *p*-value cutoff of 0.05.

### Data accessibility

All ATAC-seq and RNA-seq data generated in this study has been submitted to the National Center for Biotechnology Information (NCBI) Sequence Read Archive (SRA) under project ID PRJNA960830. All other raw data is available upon reasonable request.

### Data visualization and schematics

Heatmaps were generated in R using ComplexHeatmap^159^ 2.20.0. Unless otherwise noted, all other plots were generated in R using ggplot2^160^ 3.5.1. Schematics were created using BioRender.com.

## Acknowledgements

The authors would like to acknowledge Dr. Andrew Ewald for comments on the manuscript, Dr. WanYing “Jenny” Lin for technical advice, and Mr. James Forsmo for feedback on experimental designs and interpretation of results. This work was supported by a Physician Scientist Training Program Microgrant (J.S.H.), a Sol Goldman Pancreatic Cancer Pilot Project Grant (J.S.H and R.K.), the National Institutes of Health (NIH) (R01HG012357, R.K.) and the David & Lucile Packard Foundation (2020-71380, R.K.). J.S.H is supported by a McGlothlin Fellow-to-Faculty Award, a Conquer Cancer Foundation Young Investigator Award, and a T32 Training Grant (5T32CA009071) from the NIH. R.K.D is supported by an NSF Graduate Research Fellowship. Parts of this work were carried out at the Advanced Research Computing at Hopkins (ARCH) core facility, which is supported by the National Science Foundation (NSF) grant number OAC 1920103. FFPE processing, sectioning, and H&E staining was carried out by the Johns Hopkins Oncology Tissue Services (OTS) core facility. ATAC-seq libraries were prepared and sequenced in the Johns Hopkins Single Cell and Transcriptomics core facility. FACS was carried out at the Wilmer Eye Institute (EY001765) and at the Center for Cell Dynamics, both at Johns Hopkins. Dr. Lei Zheng generously provided the KPC cell lines utilized in this study.

## Author Contributions

J.S.H. and R.K. conceived the project; J.S.H. designed and carried out experiments and performed computational analyses; Z.L. helped generate barcoded lines; R.K.D. assisted with mouse surgeries; E.J.F. and H.G. provided critical feedback on bioinformatic analyses; J.S.H. and R.K. interpreted the data and wrote the manuscript; all authors commented on the manuscript; R.K. supervised the project.

## Competing Financial Interests

J.S.H. and R.K. are listed as co-inventors on a provisional patent on the use of this study’s findings for clinical metastasis risk prediction. E.J.F serves on the Scientific Advisory Board of Resistance Bio, as a consultant for Mestag Therapeutics and Merck, and receives research funding from Abbvie Inc and Roche/Genetech outside the scope of this work.

---

**Supplementary Figure 1.**
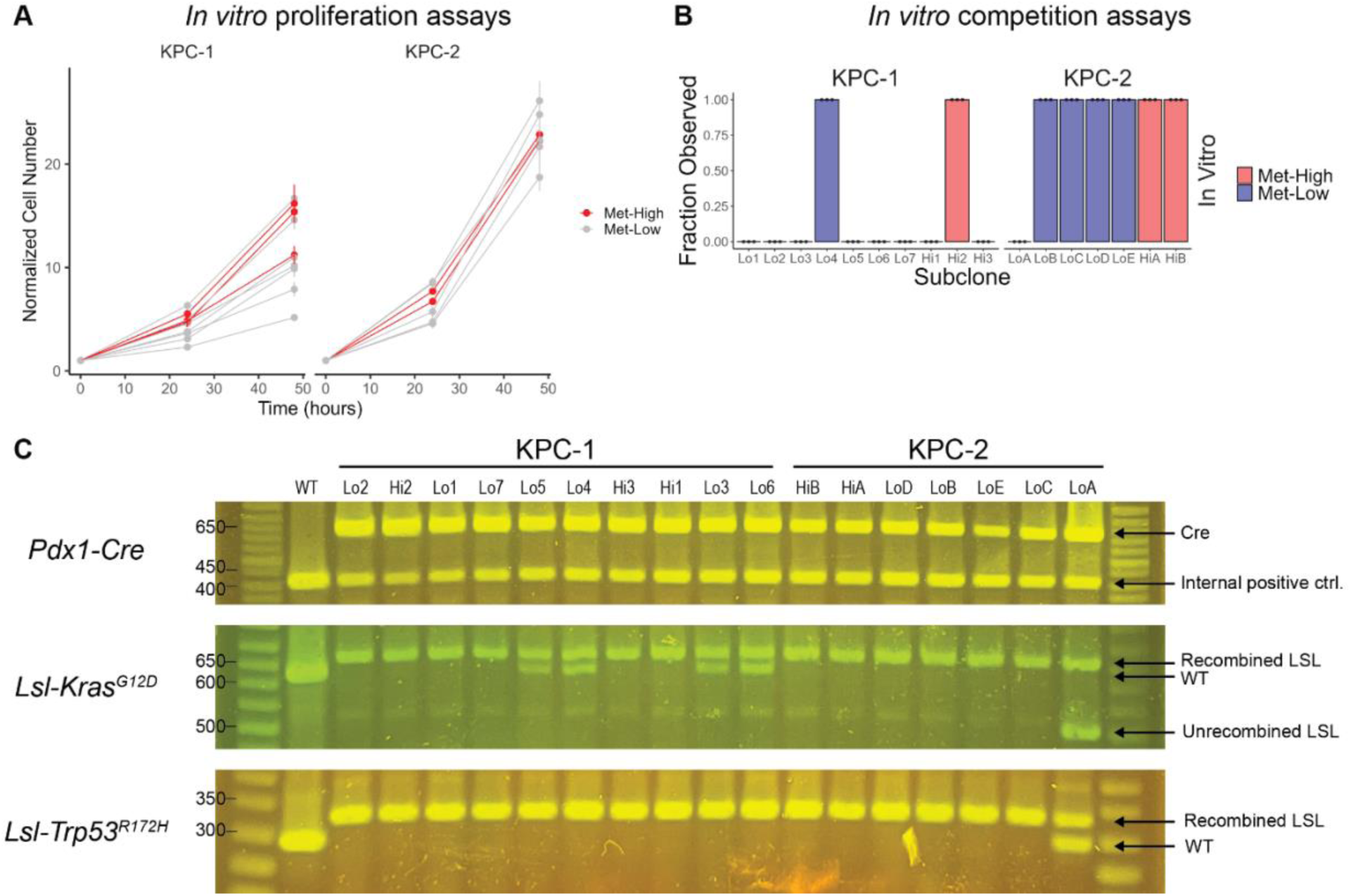
Related to Figure 1: Isolation of primary PDAC subclones with high and low metastatic potential. (**A**) *In vitro* growth curves for KPC-1- and KPC-2-derived barcoded monoclonal lines. Points represent the mean average of four technical replicates with error bars representing the S.E.M. (**B**) Fraction of three 10-cm dishes in which each subclone was observed after 28 days of passaging. (**C**) Agarose gel electrophoresis of PCR products from genotyping assays for the presence of the Pdx1-Cre transgene and for recombination of the *Lsl-Kras^G12D^* and *Lsl-Trp^R^*^172^*^H^* alleles.

**Supplemental Figure 2.**
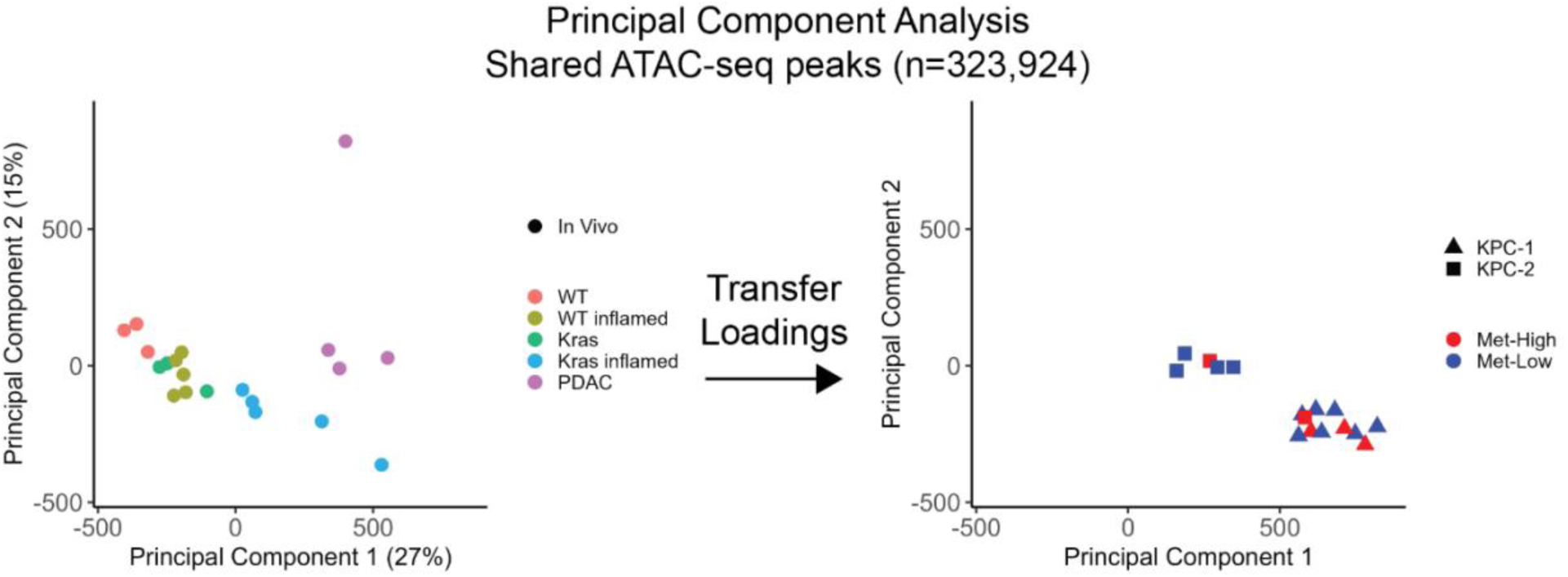
Related to Figure 2: The metastatic potential axis is orthogonal to the normal-to-PDAC and classical-to-basal axes. Principal component analysis (PCA) of scaled normalized accessibility of a consensus ATAC-seq peak set. The **left panel** shows PCA considering the WT, Kras, WT_inflamed, Kras_inflamed, and PDAC samples only. The **right panel** compares the metastasis-low and metastasis-high subclones on the same PCA axes as the left panel.

**Supplementary Figure 3.**
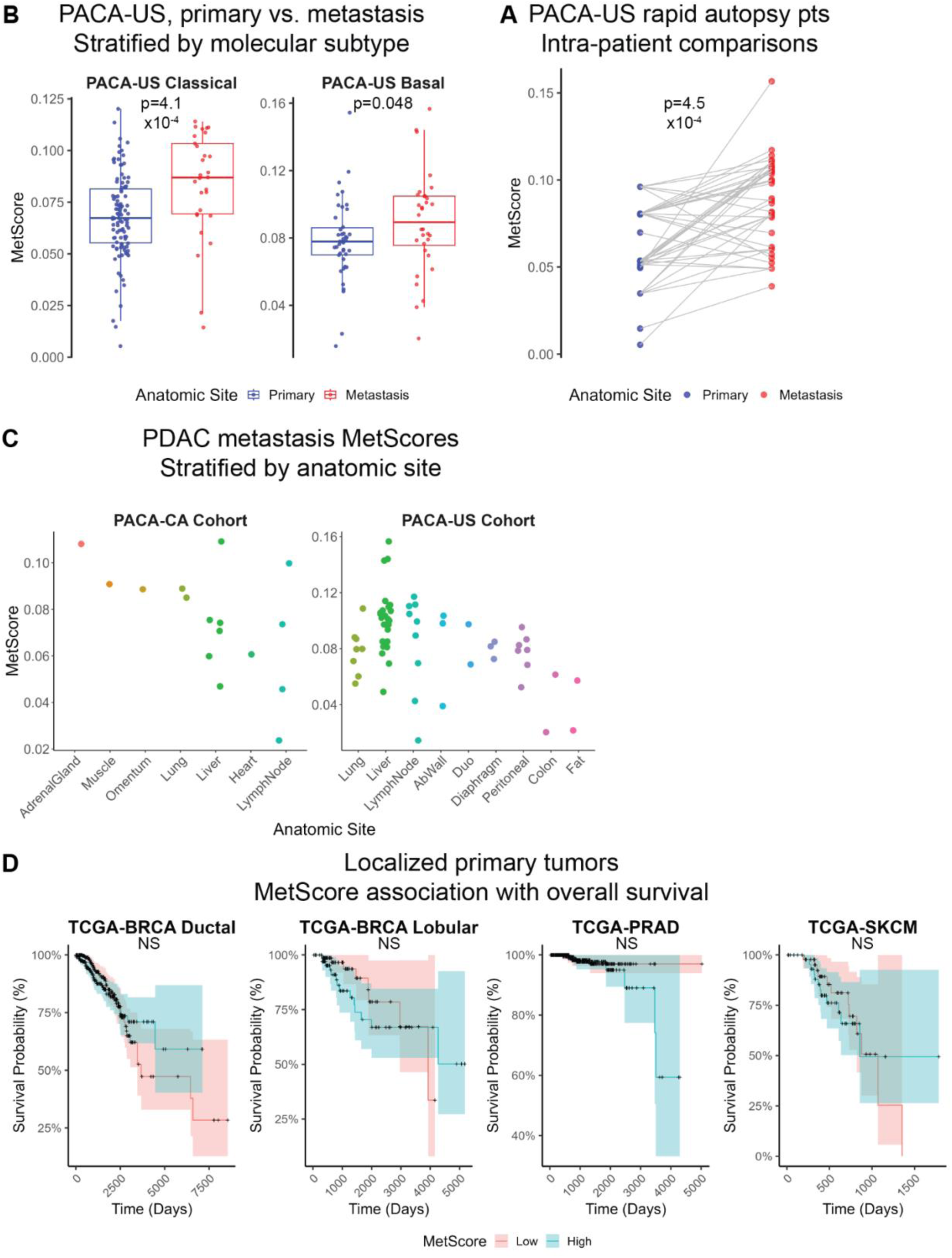
Related to Figure 7: Metastasis-high and metastasis-low genes define a metastasis signature in human carcinomas. (**A**) Box plots depicting distributions of MetScores with overlaid points representing individual tumors for primary and metastatic samples in the PACA-US cohort stratified by molecular subtype as previously annotated by the authors of the original study (Moffitt et al., 2015). *p*-values calculated using two-sided Wilcoxon rank sum tests. (**A**) Dot plots depicting MetScores for primary and metastatic samples amongst the rapid autopsy patients in the PACA-US cohort that had both primary tumors and metastases at the time of sampling. Within each patient, lines connect each primary tumor sample to each metastatic sample. *p*-value calculated using a linear mixed effect model with anatomic site as a fixed effect and patient as a random effect. (**C**) Swarm plot depicting MetScores for all metastatic samples in the indicated cohorts stratified by anatomic site. Each point represents a tumor. (**D**) Kaplan-Meier curves depicting overall survival for patients in the noted datasets stratified by MetScore. Patients in each dataset were ranked by MetScore, the top half of which being considered “High” and the bottom half considered “Low”. *p*-values were calculated using log-rank tests.

## Notes

### Competing Interest Statement

J.H. and R.K. are listed as co-inventors on a provisional patent on the use of this studies finding for clinical diagnoses. E.J.F serves on the Scientific Advisory Board of Resistance Bio, as a consultant for Mestag Therapeutics and Merck, and receives research funding from Abbvie Inc and Roche/Genetech outside the scope of this work.

